# Unremodeled plasma membrane GPI-anchored proteins trigger aberrant clathrin-mediated endocytosis

**DOI:** 10.1101/2024.07.12.603263

**Authors:** Li Chen, David K. Banfield

## Abstract

The plasma membrane has a complex organization that includes the polarized distribution of membrane proteins as well as lipids. Glycosylphosphatidylinositol-anchored proteins (GPI-AP) are ubiquitously expressed in eukaryotes and represent a functionally diverse group of proteins some of which are critical for the organization and function of the plasma membrane. Here we investigated how the plasma membrane of yeast cells responded to the accumulation of GPI-APs in which phosphoethanolamine (EtNP) had not been removed from mannose 2 (Man2) of the GPI moiety. The persistence of EtNP on Man2 prevented cleavage of a subset of GPI-APs, but the proteins were not endocytosed. Man2 unremodeled GPI-APs increased lipid disorder and generated a stress response whereby abnormal ubiquitin- and clathrin-dependent endocytosis was triggered. The resulting stress-induced endocytosis disrupted the trafficking repertoire of a subset of plasma membrane proteins. These proteins were redirected, via the multivesicular body, to numerous small vacuoles for degradation. Our findings highlight the critical importance GPI-AP Man2 remodeling for maintaining the integrity and homeostasis of the plasma membrane.

## Introduction

The plasma membrane serves as a semi-permeable barrier between the extracellular and intracellular environments of the cell. Much is known about the role of cytoskeletal components, membrane proteins, and lipids in establishing a dynamically responsive plasma membrane. Moreover, it is well understood that the interaction of certain proteins and lipids can lead to the formation of domains in the plasma membrane that are critical for the generation of signalling platforms (Simons and Toomre, 2000; Laude and Prior, 2004). Signalling platforms allow cells to respond to certain stimuli, including alterations in osmotic pressure, external pH, and the effects of cellular aging (Simons and Toomre, 2000; Parton and Hancock, 2004; Galluzzi et al., 2018). In some instances, the response to a particular stress signal can elicit a transient arrest in cell growth (de Nadal and Posas, 2022; Tognetti et al., 2020; Jiménez et al; 2020).

Glycosylphosphatidylinositol-anchored proteins (GPI-APs) are an evolutionarily conserved class of eukaryotic membrane proteins assembled in the endoplasmic reticulum (ER) via the *en bloc* addition of a lipid moiety following removal of a signal sequence (Kinoshita. 2016; Kinoshita, 2020; Muñiz and Zurzolo, 2014; Paladino et al., 2015). The GPI anchor of nascent GPI-APs undergoes extensive remodeling prior to the transport of these proteins from the ER to the plasma membrane via the Golgi (Kinoshita, 2020; Muñiz and Zurzolo, 2014). While GPI-APs are ubiquitous in eukaryotic cells their functions can vary tremendously and include roles in cell-cell adhesion, cell wall biosynthesis, signal transduction, the immune response, in the organization of lipids in the plasma membrane, and as enzymes and receptors (Laude and Prior, 2004; Paladino et al., 2015; Simons and Toomre, 2000).

The GPI-anchor is comprised of a glycan core that consists of three mannose residues, one of which is attached to the protein (Man 3) and the other two are modified through the attachment of phosphoethanolamine (EtNP). Gpi7p (PIG-V in human) transfers EtNP to Man2 of GPI-APs in the ER (Benachour et al., 1999). Ted1p (PGAP5 in human) in the ER and Dcr2p in the Golgi remove the EtNP on Man2 before the GPI-APs reach to the PM (Fujita et al., 2009; Manzano-Lopez et al., 2015; Chen et al., 2021). The removal of EtNP from Man2 is an evolutionarily conserved remodeling event whereas the removal of EtNP from Man1 has not yet been observed on human GPI-APs (Kinoshita, 2020). Glucosamine on Man1 is linked to phosphatidylinositol (PI) which is in turn modified by the addition and remodeling of various lipids (Kinoshita, 2016; Kinoshita, 2020). Most GPI anchor remodeling events occur in the ER, and they are often a prerequisite for the robust export of GPI-APs from this organelle. In metazoans, some GPI-APs are cleaved and thereafter are released from the extra-cytoplasmic side of the plasma membrane (Müller, 2018; Fujihara and Ikawa, 2016; Müller and Müller, 2023), while others, such as the folate receptor, are endocytosed. In contrast, numerous GPI-APs in budding yeast cells are cleaved once they reach the cell surface, whereupon the protein becomes a constituent of the cell wall (Kitagaki et al, 2002; Vogt et al., 2020; Yin et al., 2007). Relatively little is understood about features of the GPI moiety that are required for their cleavage from the protein, nor it is understood how remodeling defects impact the function of GPI-APs that reach the plasma membrane (Müller, 2018). Nevertheless, GPI-AP remodeling defects and /or deficiencies in the cleavage of the GPI moiety from the proteins are likely to impact plasma membrane homeostasis and have far reaching physiological consequences (Fujihara and Ikawa, 2016, Paladino et al., 2015; Sevcsik et al., 2015).

We have previously reported that incompletely remodeled GPI-APs elicit stress responses that include activation of the cell wall integrity pathway and non-canonical activation of the spindle assembly checkpoint (Chen et al., 2021). Here, we report that a failure to remove EtNP from Man2 of yeast GPI-APs renders them poor substrates for cleavage. Moreover, we show that at least one unremodeled, uncleaved GPI-AP is not endocytosed. Rather, the persistence of uncleaved, incompletely remodeled GPI-APs in the plasma membrane increased the lipid disordered phase of membranes and triggers abnormal ubiquitin- and clathrin-dependent endocytosis of select proteins, which are then degraded in an ESCRT-dependent manner in the vacuole.

We conclude that the accumulation of uncleaved, Man2 unremodeled GPI-APs disrupts plasma membrane homeostasis, triggering aberrant endocytosis in response. Our findings highlight the critical importance GPI-AP Man2 remodeling for maintaining the integrity and homeostasis of the plasma membrane. The identification of abnormal clathrin-mediated endocytosis as a response to such perturbations suggests a novel means by which plasma membrane stress signals are transmitted to the interior of the cell.

## Results

### GPI-anchored proteins (GPI-APs) with ethanolamine phosphate (EtNP) attached to mannose-2 (Man2) exhibit altered distributions in whole-cell detergent extracts and induce increased lipid disorder in cells

Our previous study (Chen et al., 2021) demonstrated that GPI-APs bearing EtNP on Man2 could still be delivered to the plasma membrane, and that their presence there induced a stress signal that triggered non-canonical activation of the spindle assembly checkpoint (SAC). From these findings, we hypothesized that GPI-APs containing EtNP on Man2 disrupt plasma membrane homeostasis in some manner that generates a stress response and concomitant arrest of cell growth.

To address the nature of the prospective cellular perturbation, we initially considered the possibility that Man2 unremodeled GPI-APs might display altered biochemical properties which impact the functional integrity of the plasma membrane. A commonly used method to examine the characteristics of membrane proteins is to monitor their partitioning in detergent extracts (Lingwood and Simons, 2007). When detergent extraction is conducted at 4°C, certain integral membrane proteins including GPI-APs, are found predominantly in detergent-resistant membranes (DRMs) (Bagnat et al., 2000; Bagnat et al., 2001).

To investigate the properties of Man2 unremodeled GPI-APs in detergent extraction experiments, we used a yeast strain in which EtNP was permanently added to Man2, hereafter referred to as the IPEM2 strain ((induced permanent phosphoethanolamine on **<u>m</u>**annose **<u>2;</u>** Chen and Banfield, 2022). IPEM2 cells can grow in the presence of glucose (conditions hereafter denoted as IPEM2-Glu) as the expression of *GPI7* (which encodes the sole enzyme that adds EtNP to Man2 in budding yeast cells) is suppressed. However, when IPEM2 cells were grown in galactose/raffinose containing media (conditions hereafter denoted as IPEM2-GR,) EtNP is added to Man2 but cannot be removed as this strain lacks the genes encoding the enzymes that remove EtNP from Man2 of GPI-APs (i.e., *TED1* and *DCR2*; Chen et al, 2021; Chen and Banfield, 2022). Figure 1A depicts the commonly used strains in this study and their corresponding genotypes. Importantly, although permanent EtNP attachment to Man2 generates a lethal phenotype eventually, approximately 80% of IPEM2 cells remained viable over the 7-hour assay period used the experiments we describe in this study (Figure 1A and 1B).

**Figure 1.**
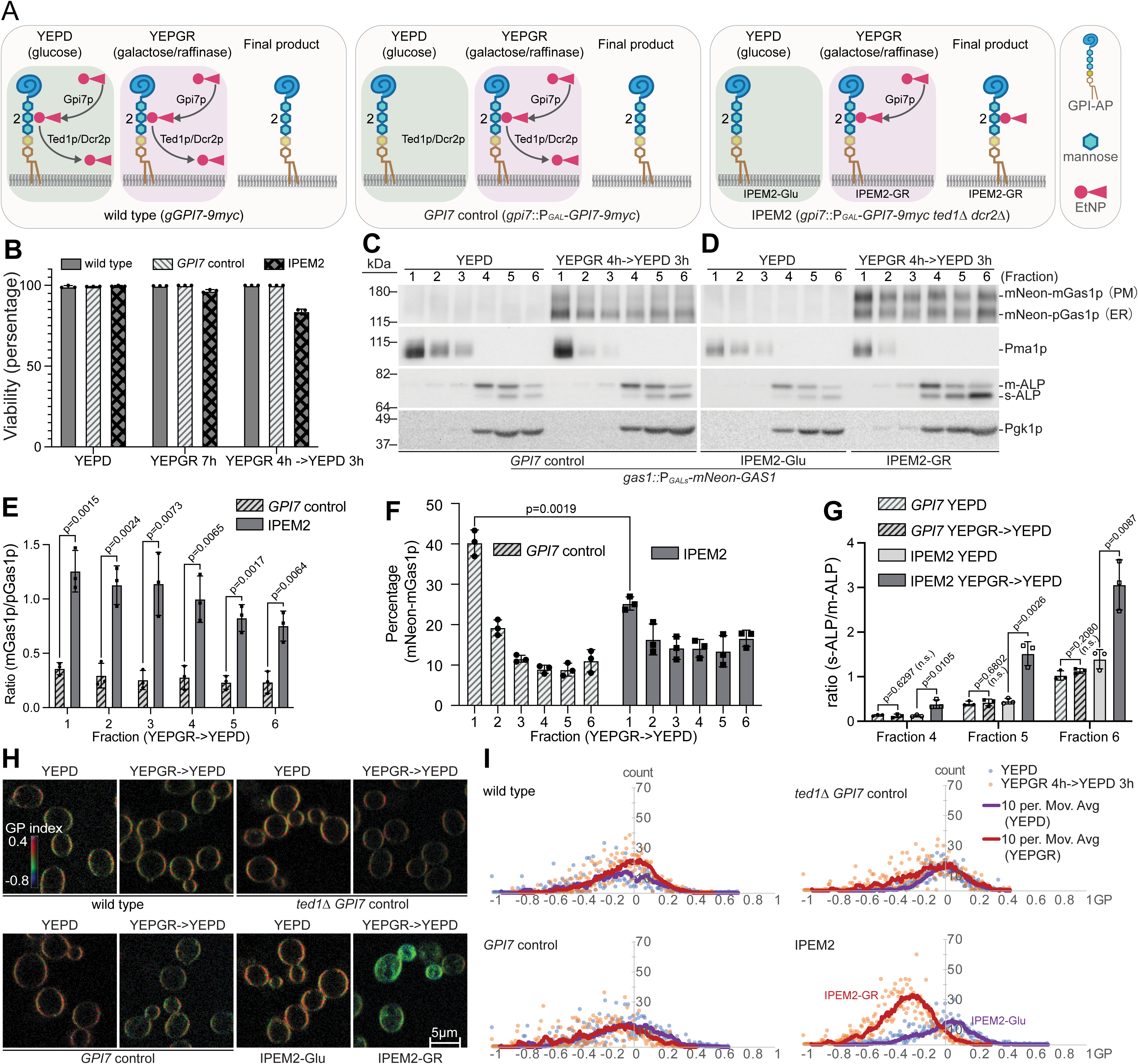
Man2 unremodeled GPI-APs display altered biochemical properties and increase the lipid disordered phase of membranes. (A) Schematic representations and genotype annotations of the commonly used yeast strains in this study. (B) More than 80% of IPEM2 cells remain viable 7 hours after induction of Gpi7p synthesis. Yeast strains were grown under the indicated growth conditions and thereafter labelled by using a LIVE/DEAD Yeast Viability Kit. More than 100 cells from each biological repeat (n=3) of were scored and the average number of viable cells plus the standard deviation (SD) are presented. (C) Protein partitioning profile of select proteins from cells expressing a galactose-inducible / glucose-repressible GPI-AP reporter protein (mNeon-Gas1p) as well as the enzyme responsible for the addition of EtNP to Man2 of GPI-APs (Gpi7p). *GPI7* expression is under the control of the GAL1/10 promoter whereas nNeon-Gas1p expression is under the control of an attenuated GAL1/10 promoter. Proteins were subjected detergent extraction at 4°C and thereafter processed and subjected to SDS-PAGE. See Figure1A for strain details. (D) Protein partitioning profile of select proteins from cells and with galactose-inducible / glucose-repressible expression of Gas1p and Gpi7p and lacking the GPI-AP Man2 remodelases (*TED1* and *DCR2*). Proteins were subjected detergent extraction at 4°C and thereafter processed and subjected to SDS-PAGE. See Figure1A for strain details. (C) and (D) depict one replicate of three independent replicates. Pma1p is a protein that displays resistance to extraction with detergent whilst ALP (Pho8p) and Pgk1p are vacuolar and cytoplasmic proteins (respectively) that are reportedly not resistant to extraction with detergent. (E) Quantification of the ratio of mNeon-mGas1p versus mNeon-pGas1p for each fraction from experiments depicted in (C) and (D) (average + SD). (Ratio=mNeon-mGas1p (fraction n) / mNeon-pGas1p (fraction n). Three independent detergent extraction datasets were used in the calculations and the average value plus the SD is presented. The solid bar denotes data from the yeast strain depicted in (D) and the crosshatched bar denotes data from experiments depicted in (C). (F) Quantification of the percentage of mNeon-mGas1p from each fraction from experiments depicted in panels C and D. out of the total fractions (average + SD). (Percentage = mNeon-mGas1p (fraction n) / total mNeon-mGas1p (fraction 1-6). Three independent detergent extraction datasets from cells grown in YEPGR media were used in the calculations and the average value plus the SD is presented. The solid bar denotes data from the yeast strain depicted in (D) and crosshatched bar denotes data from the experiments depicted in (C). (G) Quantification of the ratio of s-ALP versus m-ALP in fractions 4 – 6 in panels C and D. (Ratio = s-ALP (fraction n) / m-ALP (fraction n). Three independent detergent extraction datasets from cells grown in YEPGR media were used in the calculations and the average value plus the SD is presented. Data from the various strains are indicated by bars. (H) The GP values from the indicated Di-4-ANEPPDQH-stained yeast strains. The represented GP images are pseudo coloured and extend over the range indicated by the inserted colour bar. Pseudo colouring was constructed with an ImageJ macro (see Materials and Methods for further details). (I) Corresponding GP value histogram of cells presented in (E) (n=40). The moving average trendline was set to 10. The light blue/purple trace depicts GP values from cells cultured in YEPD (IPEM-Glu) whereas the orange/red trace depicts GP values from cells cultured in YEPGR media (IPEM-GR).

To monitor the trafficking of GPI-APs, we introduced a prototypical GPI-AP, Gas1p, into the IPEM2 strain as a galactose-inducible fusion to mNeon Green (mNeon-Gas1p), where the coding sequence of mNeon green was inserted into the *GAS1* gene after the sequence that encodes the signal peptide (Chen et al., 2021; Chen and Banfield, 2022). Due to the attenuated GAL1/10 promoter, the induction of mNeon-Gas1p occurs later than that of Gpi7p, ensuring that mNeon-Gas1p has EtNP attached to Man2 (Chen and Banfield, 2022). The molecular weight of ER resident Gas1p (pGas1p) differs from its plasma membrane-localized form (mGas1p) due to differences in their glycosylation profiles (Nuoffer et al., 1991). Trafficking of mNeon-Gas1p and expression of *GPI7* were monitored by immunostaining (Figure 1C and 1D, Figure S1A).

To examine the partitioning of mNeon-Gas1p, IPEM2-Glu and IPEM2-GR cells were lysed and extracted with 1% Triton X100 at 4°C for 30 mins before being loaded onto a step gradient and subjected to centrifugation. Following centrifugation, fractions were collected from the top (where DRMs fractionate) to the bottom of the gradient and processed for SDS-PAGE and immunostaining (Figure 1C and 1D).

As anticipated, in cells expressing the Man2 remodelases Ted1p and Dcr2p (i.e., the GPI7 control cells, Figure 1C) (Chen et al., 2021), the mature / fully remodeled form of mNeon-Gas1p was found predominately in DRM fractions (as judged by the co-fractionation with the canonical yeast DRM resident protein Pma1p, Figure 1C). In IPEM2-GR cells, we observed an incremental increase in the amount of mNeon-mGas1p across all fractions, which implies that EtNP-attachment to Man2 resulted in an increased portion of PM-associated mNeon-Gas1p (Figure 1D and 1E). As Gas1p is a cell wall associated protein (Yin et al., 2005; Yin et al., 2007) this finding suggests that Man2 unremodeled Gas1p is a poor substrate for cleavage and hence association of the protein with the cell wall.

The percentage of mNeon-mGas1p in fraction 1 represents approximately 40% of total mNeon-mGas1p (across fractions 1 - 6) in *GPI7* control (Figure 1C and 1F); whereas in the IPEM2-GR cells the percentage of the mGas1p protein from the corresponding fraction (i.e. fraction 1) was reduced to approximately 25% of the total (across fractions 1 – 6, Figure 1F). Nonetheless, the partitioning of Pma1p, Pho8p and Pgk1p (Figure 1D) was similar across all strains and growth conditions tested (Figure 1C and 1D). In sum, we therefore concluded that Gas1p containing EtNP on Man2 displayed altered biochemical characteristics that distinguished it from the completely remodeled form of the protein.

ALP (the vacuolar resident alkaline phosphatase encoded by *PHO8*) and the cytosolic protein Pgk1p were both present in the non-DRM fractions (lanes 3 – 6, Figure 1C), as expected. ALP is synthesized as an enzymatically inactive precursor that is activated in the vacuole by limited proteolysis (creating m-ALP) but nevertheless remains a vacuolar transmembrane protein (Klionsky and Emr, 1989), further processing results in the release of m-ALP from the vacuolar membrane, generating a luminal, unanchored form of the protein (designated s-ALP). The ratio of s-ALP over m-ALP in each of the non-DRM fractions from IPEM2-GR cells was significantly increased (Figure 1G), suggesting that the vacuoles of IPEM2-GR cells have may have altered protease activity. Based on these DRM extraction experiments it appears that Gas1p (and by inference other GPI-APs) in which EtNP is added to Man2 but not removed, may also have altered biochemical properties. However, incompletely remodeled Gas1p did not impact the partitioning of at least one other DRM resident protein (i.e., Pma1p, Figure 1C and 1D).

The findings presented in Figure 1 (panels B – G) prompted us to explore whether Man2 unremodeled GPI-APs affected lipid homeostasis. The plasma membrane is comprised of lipid domains that either display a more ordered state (L_O_) enriched in sterols and sphingolipids, or a more disordered state (L_D_) enriched in lipids containing unsaturated fatty acids and exhibit greater mobility of lipids (Sezgin et al., 2017). Maintaining the balance of L_O_ versus L_D_ domains is thought to play a central role in establishing the functional landscape of the plasma membrane (Simons and Toomre, 2000; Laude and Prior, 2004). To address the prospect that GPI-APs in IPEM2-GR cells might perturb the ratio of L_O_/L_D_ domains, we employed the aminonaphthylethenylpyridinium voltage-sensitive dye di-4-ANEPPDHQ in quantitative confocal fluorescence microscopy experiments (Owen et al., 2011). This dye is excited at 488 nm yet results in a peak emission wavelength of ∼560 nm in the lipid ordered phase and ∼610 nm in the disordered phase. By measuring the fluorescent intensity of the two emissions, we were able to monitor the changes in L_O_/L_D_ domains. The spectral shift of di-4-ANEPPDHQ to ∼560 nm or ∼610 nm allows calculation of the generalized polarization (GP) which is a relative but quantitative measure of lipid packing (Amaro et al., 2017; Jin et al., 2005; Zhao et al., 2015). In such experiments we observed an overall increase in the L_D_ proportion of membranes (or relatively loosely packed membranes) in IPEM2-GR, but not in IPEM2-Glu cells or control strains (Figure 1H and 1I). Thus, the biochemical properties of GPI-APs in IPEM2-GR cells are altered, and these cells display an increase in membrane L_D_.

#### GPI-APs are not cleaved or endocytosed in IPEM2-GR cells

In budding yeast cells, most GPI-APs are constituents of the cell wall, where the GPI moiety is cleaved at the plasma membrane and the protein is then covalently attached to the cell wall (Yin et al., 2005; Müller, 2018). The results of our spectral ratiometric imaging experiments (Figure 1H and 1I) revealed that membranes from IPEM2-GR cells exhibited an increase in L_D_, which likely impacts the extent to which regions of the plasma membrane cluster sterols and sphingolipids (Simons and Toomre, 2000; Laude and Prior, 2004). As GPI-APs appear to be delivered to the cell surface (Chen et al., 2021) the observed membrane perturbations in IPEM2-GR cells may be a consequence of accumulation of uncleaved proteins in the plasma membrane. To investigate this further, we examined the fate of newly synthesized Gas1p in IPEM2-GR cells. We generated an IPEM2 strain constitutively expressing mScarlet-Gas1p and a copy of mNeon-Gas1p whose expression was under the control of the GAL1/10 attenuated promoter (Figure 2A). In control strains, *de novo* synthesized mNeon-Gas1p was predominantly targeted to the incipient daughter cell, together with newly synthesized mScarlet-Gas1p. Notably, the distribution mScarlet-Gas1p was not uniform (as judged by fluorescence intensity) suggesting that protein from the daughter cell was not free to diffuse (asterisk, Figure 2A). However, in IPEM2-GR cells, both mNeon-Gas1p and mScarlet-Gas1p were uniformly distributed around the cell periphery. This observation is consistent with the data from the DRM experiments (Figure 1D), and suggest that in IPEM2 cells, Gas1p cannot be cleaved and cross-linked to the cell wall, and thus the protein is free to diffuse in the plasma membrane (Figure 2A).

**Figure 2.**
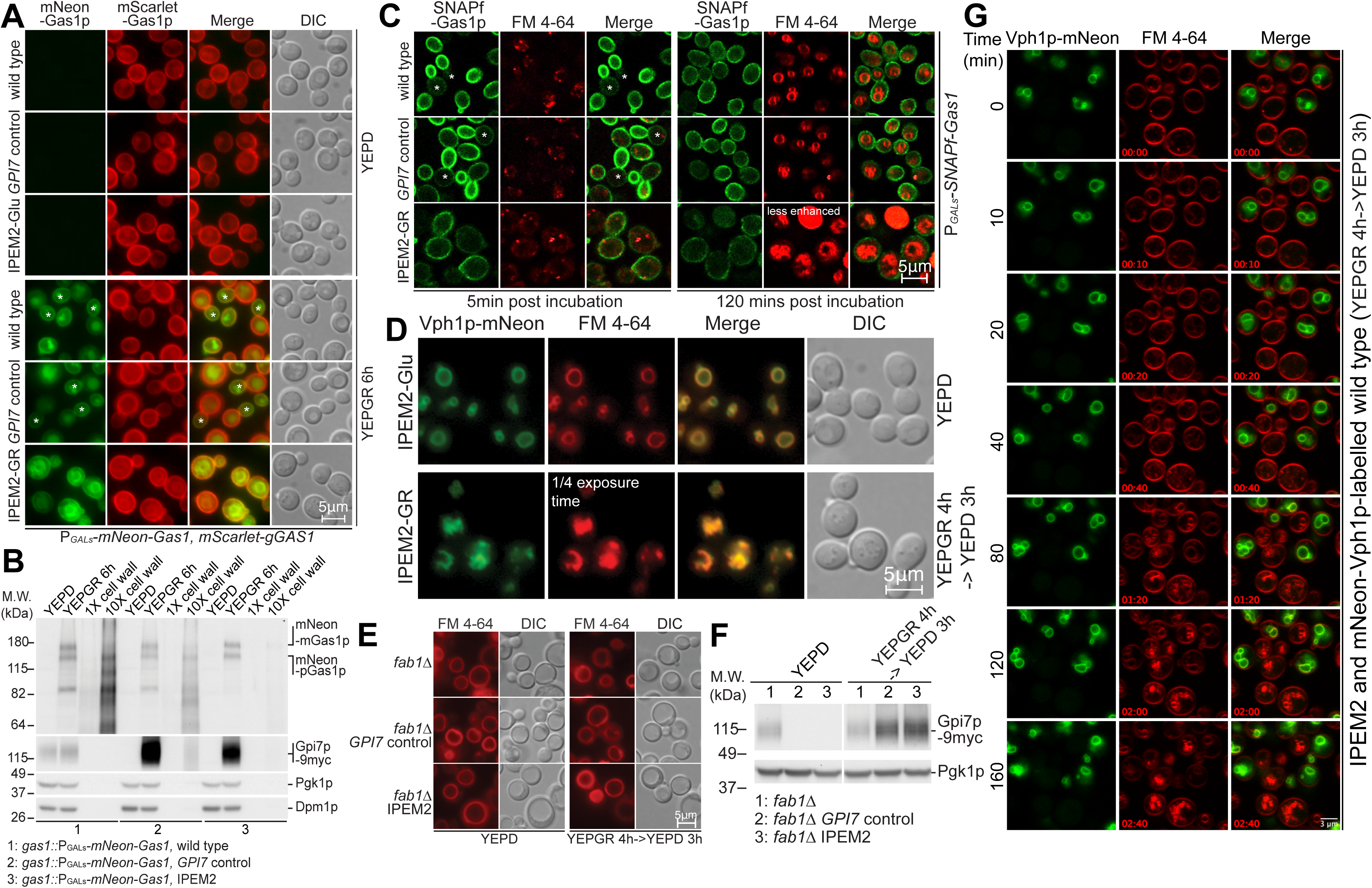
The prototypical GPI-AP Gas1p is not cleaved or endocytosed in IPEM2-GR cells. (A) Newly synthesized Gas1p is targeted to the nascent bud in WT and *GPI7* control cells, but uniformly distributed on the plasma membrane in IPEM-GR cells. The indicated strains were grown in either YEPD or in YEPGR containing media for 6 hours to induce the synthesis of mNeon-Gas1p and Gpi7p. The asterisks indicate the location of newly synthesized mNeon-Gas1p in WT and *GPI7* control cells. (B) mNeon-Gas1p IPEM2-GR cells is not associated with the cell wall. Yeast strains were grown as indicated and treated as described in the Materials and Methods to separate the cell wall from other cellular constituents. Pgk1p and Dpm1p and Gpi7p serve as markers for the cytoplasmic and ER membrane fractions. mNeon-Gas1p was detected using an anti-Gas1p antibody. (C) mNeon-Gas1p from IPEM2-GR cells reaches the plasma membrane but is not endocytosed. The indicated strains expressing GAL1/10 promoter driven SNAP-tagged Gas1p were labelled with SNAP-Surface 488 and the fate of the labelled protein was assessed 5 minutes and 120 minutes after removal of excess dyes and galactose + raffinose. The asterisks indicate the exclusion of *de novo* synthesized plasma membrane localized SNAP-Gas1p from the mother cell in WT and *GPI7* control cells. (D) The vacuolar resident integral membrane protein Vph1p-mNeon localizes to the FM 4-64 positive membrane structures in IPEM2-GR cells. (E) The membranous structures that accumulate in IPEM2-GR cells (in panels C and D) represent numerous small vacuoles. Note that deletion of *FAB1* in the IPEM2 strain generates a single large vacuole. (F) Expression profile of Gpi7p in the yeast strains depicted in (E). The indicated strains were grown in YEGR for 4 hours and subsequently for 3 hours in YEPD. Proteins samples were thereafter collected and subjected to SDS-PAGE and immunostaining. *GPI7* is driven by the GAL1/10 promoter in *fab111 GPI7* and *fab111* IPEM2 cells, and by *GPI7*’s endogenous promoter in *fab111* cells. Pgk1p serves as a gel loading control. (G) The kinetics of FM4-64 uptake and transport are indistinguishable in IPEM2-GR and WT cells. WT cells expressing Vph1p-mNeon were mixed with IPEM2-GR cells (as indicated) and thereafter cells were labelled with FM4-64 dye. Cells were imaged over the time course shown. Note that only IPEM2 cells show increased FM4-64 fluorescence intensity (from the 80-minute time point onwards).

To more conclusively address whether Man2 unremodeled Gas1p is a poor substrate for cleavage and cross-linking to the cell wall, we examined the distribution of mNeon-Gas1p in control and IPEM2 (-Glu and -GR) cells. In these cells, mNeon-Gas1p expression was controlled by an attenuated GAL1/10 promoter (GALs; Figure 2B). After inducing of mNeon-Gas1p synthesis, we prepared whole-cell and cell wall extracts for immunostaining (Figure 2B). While control strains and IPEM2-GR cells had equivalent amounts of the precursor and mature forms of mNeon-Gas1p, mNeon-Gas1p was undetectable in cell wall extracts from IPEM2-GR cells (strain 3; Figure 2B). These observations support the idea that the EtNP modification on Man2 impairs the cleavage and cross-linking of Gas1p to the cell wall. Similar findings were also obtained for the stress-induced GPI-anchored cell wall protein Sed1p (Shimoi et al., 1998; Yin et al., 2005) (Figure S2A and B). Our data agree with experiments conducted with *cdc1* mutants, which are defective in the removal of EtNP from Man1 of GPI-APs (Vazquez et al., 2014). It appears that failure to remove EtNP from Man2 interferes with cleavage of the GPI-moiety from at least two GPI-APs, and this observation may be more broadly applicable given the number of such proteins covalently linked to the yeast cell wall (Yin et al., 2005).

The observation that Gas1p is not cleaved at the plasma membrane prompted us to ask whether plasma membrane-resident Gas1p was endocytosed in IPEM2-GR cells, as this could provide a means to re-establish plasma membrane homeostasis. To monitor the fate of plasma membrane-localized Gas1p in IPEM2-GR cells, we introduced an inducible SNAP-tagged Gas1p fusion protein (SNAP-Gas1p) and used the membrane impermeable dye SNAP-Surface 488 to label only SNAP-Gas1p fusion proteins that had reached the plasma membrane (Keppler et al., 2004) (Figure 2C). After inducing of SNAP-Gas1p synthesis, we incubated cells in the presence of SNAP-Surface 488 and the styryl dye FM 4-64 (to visualize endosomes and the vacuole) (Vida and Emr, 1995). We then washed cells to remove excess dyes and galactose+raffinose and resuspended them in media containing glucose to suppress *de novo* synthesis of SNAP-Gas1p and Gpi7p. FM 4-64 positive puncta (presumably endosomes) were evident in all strains examined following 5 minutes of incubation at 25°C, and following 120 mins, the limiting membrane of the vacuole was labelled with FM 4-64 in control strains (Figure 2C). However, after 120 mins at 25°C, neither the controls nor IPEM2-GR cells showed any evidence of internalized SNAP-Gas1p (Figure 2C). In IPEM2-GR cells, *de novo* synthesized SNAP-Gas1p (5-minute time-point) was distributed equally on the surface of mother and daughter cells (Figure 2C), which is consistent with the data presented in Figure 2A and further supports our view that IPEM2-GR cells are deficient in cleaving GPI-APs.

Additionally, in IPEM2-GR cells FM 4-64 accumulated brightly fluorescent internal structures that we surmised represented small and often clustered vacuoles (Figure 2C and Movie EV1). The intensity of internal FM 4-64 fluorescence in IPEM2-GR cells at the 120-minute point relative to control strains suggests that IPEM2-GR cells have physiochemically abnormal vacuoles – as all strains were treated with the same concentration of FM4-64 dye for the same period prior to removing unincorporated dye.

To confirm the vacuolar origin of the FM 4-64 positive internal structures in IPEM2-GR cells, we introduced an integral membrane protein that localizes to the limiting membrane of the vacuole (Vph1p-mNeon; Figure 2D). As anticipated the brightly fluorescent internal structures apparent with FM 4-64 colocalized with Vph1p-mNeon suggesting that these structures are of vacuolar origin. To address whether the internal structures that appeared in IPEM2-GR cells represented clusters of small vacuoles we deleted *FAB1*. *FAB1* encodes the only 1-phosphatidylinositol-3-phosphate 5-kinase in the yeast genome and Fab1p is responsible for the synthesis of PI3,5P2 (Yamamoto et al., 1995). Cells that lack *FAB1* are defective in vacuolar fission and consequently contain a single large vacuole (Yamamoto et al., 1995). When we examined vacuole morphology in IPEM2-GR *fab1*Δ cells with FM 4-64 dye we observed a single large vacuole (Figure 2E and F). Based on this observation, as well as the colocalization of Vph1p and FM 4-64 with the internal structures, we concluded that IPEM2-GR cells contained numerous intact small vacuoles (Movie EV1) that can nevertheless coalesce to form a single larger vacuole when *FAB1* is deleted. In budding yeast cells more than 130 genes, when defective, cause a fragmented / small vacuoles phenotype (Hurst and Fratti, 2020; Michaillat and Mayer, 2013), but given the scope of this study, the cause of numerous small vacuoles in IPEM2 cells was not explored further.

To assess if the small, clustered vacuole phenotype of IPEM2-GR cells was the result of enhanced endocytosis, we used FM 4-64 to label the plasma membrane of a mixture of IPEM2-GR and mNeon-Vph1-labelled wild type cells, and simultaneously followed the endocytosis of FM4-64 into cells over the course of 160 mins using an LSM980 confocal microscope. Cells expressing mNeon-Vph1 are distinguishable from their IPEM2-GR counterparts by the presence of green vacuoles (Figure 2D and Movie EV2). Several FM 4-64 positive puncta of equal intensity were visible in both yeast strains beginning 20 mins post dye incubation. By 40 mins post FM4-64 incubation numerous FM4-64 positive puncta were apparent in IPEM2-GR cells, and fewer, less intensely fluorescent puncta where visible in mNeon-Vph1 expressing cells (Figure 2G and Movie EV2). At 80 mins post FM4-64 incubation and time points beyond this, IPEM2-GR cells contained numerous, what appeared to be, clusters of small vacuoles that were far brighter in intensity than FM4-64 labelled vacuoles from mNeon-Vph1 expressing cells (Figure 2G and Movie EV2). Based on these data, we concluded that the rate of FM4-64 uptake into IPEM2-GR cells was indistinguishable from that in mNeon-Vph1 expressing (and otherwise wild type cells). Presumably, the increased FM4-64 fluorescence intensity observed in IPEM2-GR cells reflects some alteration to vacuoles in this strain.

### IPEM2-GR cells reroute endocytosed proteins via clathrin-mediated endocytosis into numerous small vacuoles

If the stress response in IPEM2-GR cells was due to the presence of unremodeled GPI-APs on the plasma membrane, a plausible outcome of this would be for cells to remove these proteins by endocytosis, as has been described for a misfolded GPI-AP (Satpute-Krishan et al., 2014). However, the data presented in Figure 2C revealed that SNAP-Gas1p was not endocytosed in IPEM2-GR cells. Therefore, we next asked whether IPEM2-GR cells altered endocytosis more generally to achieve a correction of the plasma membrane perturbation.

To examine endocytosis in IPEM2-GR cells, we assessed the trafficking of the well-studied R-SNARE Snc1p (Lewis et al., 2000). In wild type cells, Snc1p cycles between the Golgi, the plasma membrane, and the endosome; but at steady state, the protein is predominantly localized to the plasma membrane and to nascent buds (Figure 3A) (Lewis et al., 2000). However, in IPEM2-GR cells, Snc1p was largely absent from the plasma membrane, accumulating instead in numerous small vacuoles, as judged by colocalization with FM 4-64 and the presence of “free” GFP in immunoblots from whole-cell extracts (Figure 3A and 3B). GFP is resistant to vacuole-mediated proteolysis, and therefore, the presence of “free” GFP indicates that GFP-Snc1p reached the lumen of the vacuole. The absence of *de novo* synthesized Snc1p from the plasma membrane of the daughter cell suggested that the trafficking of this protein between the endosome and plasma membrane was defective in IPEM2-GR cells (Figure 3A).

**Figure 3.**
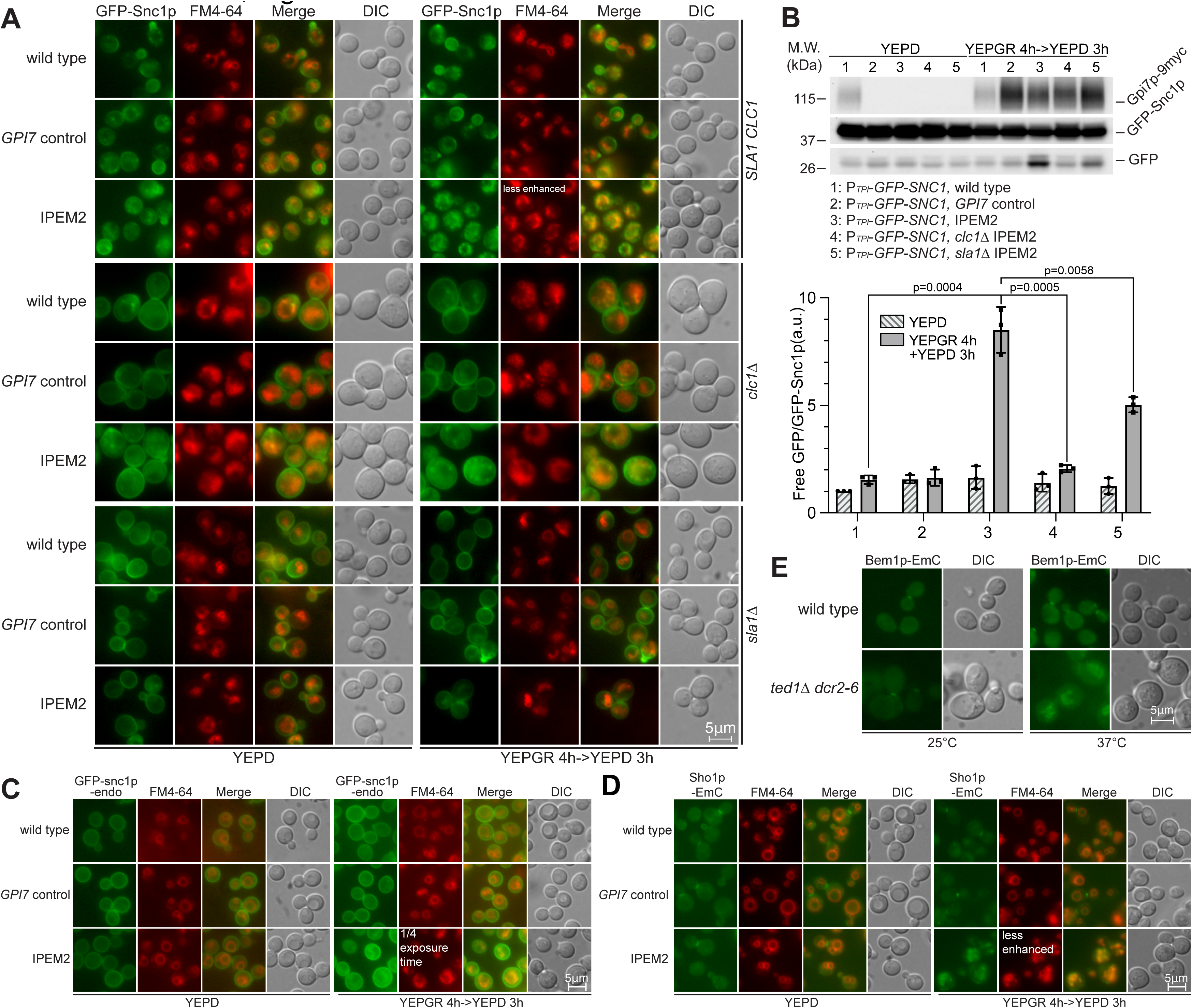
IPEM2-GR cells accumulate clathrin-mediated endocytosed proteins in numerous small vacuoles. (A) The clathrin-mediated trafficking repertoire of GFP-Snc1p is disrupted in IPEM2-GR cells. *clc111* cells are larger than their wild type counters. (B) GFP-Snc1p is re-routed to the vacuole and degraded in IPEM2-GR cells. Upper panel: representative immunoblot of GFP-Snc1p from WCEs of the various yeast strains grown under the indicated conditions. n=3; lower panel: quantification of the ratio of free GFP versus GFP-Snc1p from the various yeast strains under the indicated conditions (average + SD). n=3. a.u. denotes arbitrary units (C) In cells expressing GFP-snc1p harbouring amino acid substitutions that block clathrin-mediated endocytosis (snc1-endo) restricts the protein to the plasma membrane in IPEM2-GR cells. (D) In IPEM2-GR cells Sho1p-EmC was redirected to numerous small vacuoles. EmC denotes enhanced monomeric Citroen. (E) In *ted111 dcr2-6* cells grown at 37°C Bem1p-EmC was redirected to numerous small vacuoles. EmC denotes enhanced monomeric Citroen.

The abnormal trafficking of GFP-Snc1p in IPEM2-GR cells was clathrin-mediated, as in IPEM2-GR cells lacking the light chain of clathrin (IPEM2 *clc1ι1*) or Sla1p (an effector of clathrin-mediated endocytosis) (IPEM2 *sla1ι1*), GFP-Snc1p was uniformly distributed on the plasma membrane (Figure 3A). Given the apparent absence of free GFP in immunoblots from whole cell extracts (WCEs), and the absence of any significant colocalization of these puncta with FM 4-64 positive membranes, we concluded that GFP-Snc1p did not reach the vacuole (Figure 3A and 3B).

Further evidence for the aberrant clathrin-mediated endocytosis of GFP-Snc1p in IPEM2-GR cells was obtained by examining the localization of an endocytic mutant of Snc1p bearing substitutions of Val41 and Met43 to Ala (termed snc1p-endo; Lewis et al., 2000), which render Snc1p unable to bind a clathrin adaptor. When expressed in IPEM2-GR cells GFP-snc1p-endo was predominantly localized to the plasma membrane although some vacuolar localization was also apparent (Figure 3C). Based on these findings, we concluded that in IPEM2-GR cells that GFP-Snc1p could reach the plasma membrane, and thereafter entered the cell interior via clathrin-mediated endocytosis.

Given the observation that the polarized distribution of GFP-Snc1p on the plasma membrane was defective in IPEM2-GR cells we sought evidence of additional proteins whose distributions might also be affected. For these experiments, we chose two proteins with known roles in establishing yeast cell polarity, and in responding to stress induced by osmotic changes, termed Bem1p and Sho1p, respectively (Leeuw et al., 1995; Tatebayashi et al., 2015). Sho1p is a multi-spanning membrane protein, and in wild type cells Sho1p is found on the plasma membrane and at the bud neck (Figure 3D). Bem1p, is an SH3 domain-containing peripheral membrane protein that binds to PIP3 (Slessareva et al., 2006). In control cells and IPEM2-Glu cells Sho1p localized to the bud neck, whereas in IPEM2-GR cells Sho1p translocated to the vacuole (Figure 3D). Similarly, in *ted111 dcr2-6* cells that were cultured at 25 °C Bem1p was localized to the bud neck. However, Bem1p was relocated to the vacuole when *ted111 dcr2-6* mutant was cultured at 37 °C (Figure 3E).

The redirection of Snc1p, Bem1p and Sho1p from the plasma membrane to the vacuole in Man2 remodeling mutants was specific to the deficiency in the removal of EtNP on Man2 of GPI-APs. This was evident from the fact that deletion of the GPI-AP lipid remodeling genes *BST1* (Fujita et al., 2006), *GUP1* (Bosson et al., 2006) or *GPI7* (which adds EtNP on Man2) did not result in the mislocalization of Sho1p to vacuoles (Figure S3A and S3B). Similarly, deletion of *TED1* or *DCR2* also did not lead to mislocalization of GFP-Snc1p to the vacuole (Figure S3C and S3D). The redirection of proteins observed in IPEM2-GR cells did not apply to plasma membrane proteins generally as neither the distribution of Sur7p (a component of the eisosome; Walther et al., 2006) or Mid2p (a sensor of the cell wall integrity pathway; Rajavel et al., 1999) were altered in IPEM2-GR cells (Figure S3E and S3F). In addition, in temperature-sensitive dcw*1-3 dfg511* cells (Figure S3G), which fail to transfer the GPI-APs from the plasma membrane to the cell wall when grown at 37°C (Kitagaki et al., 2002; Kitagaki et al., 2004; Marian et al., 2020), the polarized distribution and degradation of GFP-Snc1p was not affected (Figure S3H and S3I). This suggests that it is the presence of EtNP on Man2 of GPI-APs, rather than the failure to cleave GPI-APs at the plasma membrane, that accounts for the redirection of proteins.

### Ubiquitination is a prerequisite for abnormal clathrin-mediated endocytosis in IPEM2-GR cells

Plasma membrane proteins destined for degradation are transported into the interior of the vacuole via the multivesicular body (MVB) (Babst, 2011). The formation of the MVB involves the function of a series of ESCRT complexes that act successively (Babst, 2011). To explore a role for the MVB in the degradation of GFP-Snc1p we deleted *VPS27* or *DOA4* in the IPEM2 strain. Vps27p is a component of the ESCRT-0 complex and mediates protein degradation by binding to and incorporating proteins that have been ubiquitinated into the nascent MVB (Katzmann et al., 2003), whereas Doa4p is a ubiquitin hydrolase that functions to recycle ubiquitin at a later stage in MVB biogenesis (Nikko and André, 2007).

To assess the impact of deletion of *VPS27* or *DOA4* in IPEM2-GR cells, we examined the fate of GFP-Snc1p by fluorescence microscopy and monitored the degradation of GFP-Snc1p by immunoblotting of WCE extracts. In IPEM2-GR cells, GFP-Snc1p accumulated in internal puncta as well as in vacuoles (as judged by co-localization with FM 4-64 dye) (Figure 4A). In contrast, in IPEM2-GR cells in with either *VPS27* or *DOA4* deleted (IPEM2-GR *vps27Δ* and IPEM2-GR *doa411)*, the majority of GFP-Snc1p was found in internal structures that did not colocalize with FM 4-64 dye (Figure 4A and 4B). Consistent with the fluorescence microscopy data, immunoblotting of WCEs revealed that when either *VPS27* or *DOA4* were deleted from IPEM2-GR cells (IPEM2 *vps27Δ* and IPEM2 *doa4Δ*), less protease resistant GFP was evident, compared to IPEM2-GR cells in which these genes had not been deleted (Figure 4C). These data suggest that disrupting the formation of MVBs in IPEM2-GR cells prevented delivery of GFP-Snc1p to the vacuole.

**Figure 4.**
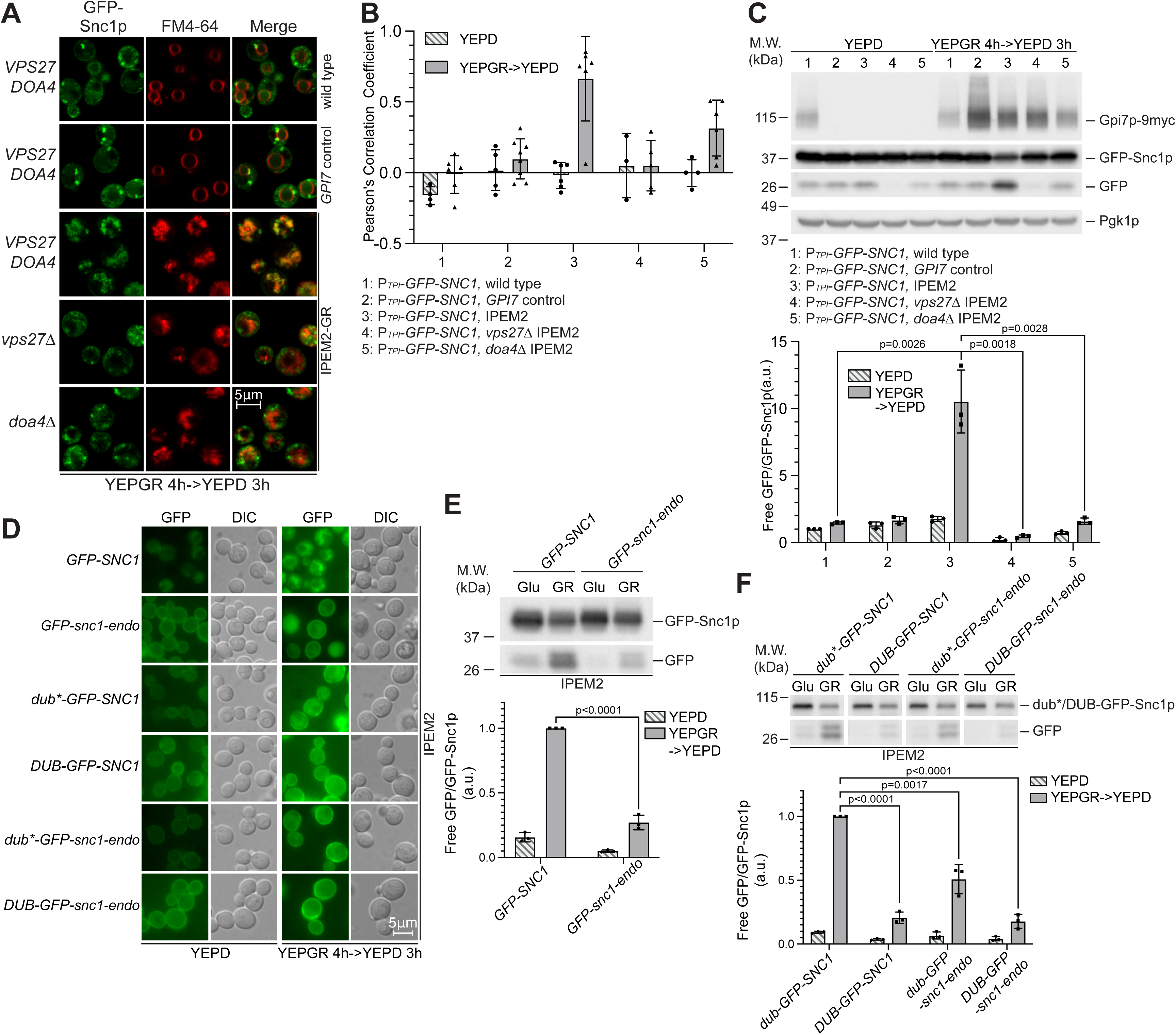
Ubiquitination is a prerequisite for the abnormal clathrin-mediated endocytosis observed in IPEM2-GR cells. (A) GFP-Snc1p is rerouted numerous small vacuoles in IPEM2-GR cells via the MVB pathway. (B) Colocalization of GFP-Snc1p puncta with FM 4-64 labelled membranes from the experiment depicted in (A) n=3-8 cells (depending growth conditions and the particular strain used). See Materials and Methods for details. (C) A functional ESCRT pathway is required for delivery of GFP-Snc1p to the vacuole in IPEM2-GR cells. Upper panel: representative immunoblotting of GFP-Snc1p from the indicated strains under the indicated conditions. n=3; lower panel: quantification of the ratio of free GFP versus GFP-Snc1p (average + SD). n=3. a.u. denotes arbitrary units. Pgk1p serves as a gel loading control. (D) The abnormal clathrin-mediated endocytosis of GFP-Snc1p requires the protein be ubiquitinated. snc1-endo denotes the endocytosis deficient mutant of Snc1p, DUB denotes the de-ubiquitination domain from Ubp7p and dub* the de-ubiquitination domain deficient mutant. See the main text for a more detailed description. (E) The clathrin-mediated endocytosis deficient mutant of GFP-Snc1p (GFP-snc1-endo) is a poor substrate for delivery to the vacuole in IPEM2-GR cells. Glu denotes growth media containing 2% glucose, Gal denotes growth media containing 2% galactose and 1% raffinose. a.u. denotes arbitrary units. Upper panel: representative immunoblotting of GFP-Snc1p under indicated conditions. n=3; lower panel: quantification of the ratio of free GFP versus GFP-Snc1p (average + SD). n=3. a.u. denotes arbitrary units. (F) Ubiquitination of GFP-Snc1p is a prerequisite for the protein’s delivery to the vacuole in IPEM2-GR cells. Upper panel: representative immunoblotting of GFP-Snc1p under indicated conditions. n=3; lower panel: quantification of the ratio of free GFP versus GFP-Snc1p (average + SD). n=3. a.u. denotes arbitrary units. Glu denotes growth media containing 2% glucose, Gal denotes growth media containing 2% galactose and 1% raffinose.

To more directly assess the role of ubiquitin in the endocytosis of Snc1p, we fused the de-ubiquitination domain of Ubp7p or the de-ubiquitination deficient domain (ubp7p; C618S) to GFP-Snc1p, generating DUB-GFP-Snc1p and dub*-GFP-Snc1p, respectively. Similar replacements were also made to the endocytosis-deficient form of Snc1p (snc1-endo) generating DUB-GFP-snc1-endo and dub*-GFP-snc1-endo (Figure 4) (Stringer and Piper, 2011). The GFP-Snc1p fusion proteins were introduced into IPEM2-GR cells, and the fate of these fusion proteins was examined by fluorescence microscopy and immunoblotting (Figure 4D, 4E, and 4F, respectively). As expected, DUB-GFP-Snc1p was uniformly distributed on the plasma membrane, and comparatively little protease-resistant GFP was apparent in WCEs from IPEM2-GR cells - data that are consistent with ubiquitination being a prerequisite for the endocytosis of Snc1p (Figure 4D and 4F). In contrast, dub*-GFP-Snc1p localized to both the plasma membrane and internal structures, and comparatively more protease-resistant GFP (∼75% more) was apparent in WCEs from these cells compared to DUB-GFP-Snc1p (Figure 4D and 4F). Although GFP-snc1-endo was a poor substrate for clathrin-mediated endocytosis (Figure 3C and Figure 4D), endocytosis still required that the protein be ubiquitinated as judged by fluorescence microscopy and immunoblots of WCEs (Figure 4D and 4F). Similar findings were also obtained with Sho1p-EmC-DUB in IPEM2-GR cells (Figure S4). In IPEM2-Glu cells, Sho1p-EmC was primarily found at the mother / daughter cell junction (Figure S4A and S4B), whereas Sho1p-EmC localized to highly fragmented vacuoles in IPEM2-GR cells (Figure S4A and S4B). Sho1p-EmC-DUB was no longer found at the mother / daughter cell junction in control cells (grown in glucose or galactose+raffinose) or in IPEM2-Glu cells (Figure S4C and S4D) Instead, Sho1p-EmC-DUB was distributed non-uniformly on the plasma membrane, revealing a role for ubiquitination in the redirection of Sho1p to the mother / daughter cell junction (Figure S4C). In IPEM2-GR cells, Sho1p-EmC-DUB was found on the plasma membrane (although the protein appeared to be largely excluded from the daughter cell plasma membrane) as well as in the numerous small vacuoles of both the mother and daughter cells (Figure S4C).

### The E3 ligase Rsp5p plays multiple roles in the clathrin-mediated endocytosis of proteins in IPEM2-GR cells

*RSP5* encodes a NEDD4 family E3 ubiquitin ligase implicated in multiple processes including the regulation of endocytosis and the sorting of proteins into the MVB (Katzmann et al., 2004). To examine the role of Rsp5p in the ubiquitin-dependent endocytosis of Sho1p, we introduced a temperature-sensitive allele of *RSP5* (*rsp5-1*) (Dunn and Hicke, 2001; Katzmann et al., 2003) into *ted1Δ dcr2-6* cells. While *rsp5-1* did not affect the localization of Sho1p-EmC grown at 37°C in otherwise wild type cells, in *rsp5-1 ted111 dcr2-6* cells grown at 37°C, Sho1p-EmC localized to the limiting membrane of the vacuole (Figure 5A) and was excluded from the vacuolar lumen, as evidenced by the absence of free GFP in immunoblots of WCEs from these cells (Figure 5B). This indicates that Rsp5p activity is not required for the initial stages of endocytosis of Sho1p in the temperature sensitive Man2 remodeling mutant, but rather is necessary for the internalization, and subsequent degradation of Sho1p in the vacuolar lumen. In contrast to Sho1p, Snc1p required Rsp5p activity for both the initial stages of endocytosis as well as for the internalization and degradation of Snc1p in the vacuole (Ma and Bird, 2019). In IPEM2-GR *rsp5-1* cells, Snc1p was still observable on the plasma membrane, and comparatively less GFP-Snc1p was delivered to the lumen of the vacuole as indicated by a reduction in the amount of “free” GFP in immunoblots of WCEs (Figure 5C and 5D). Interestingly, introduction of the *rsp5-1* allele into *ted111 dcr2-6* cells substantially reduced the numerous small vacuole phenotype of these cells at 37°C (FM4-64 staining; Figure 5A and 5C) suggesting that this phenotype arises from an imbalance in the fusion and fission of vacuolar membranes.

**Figure 5.**
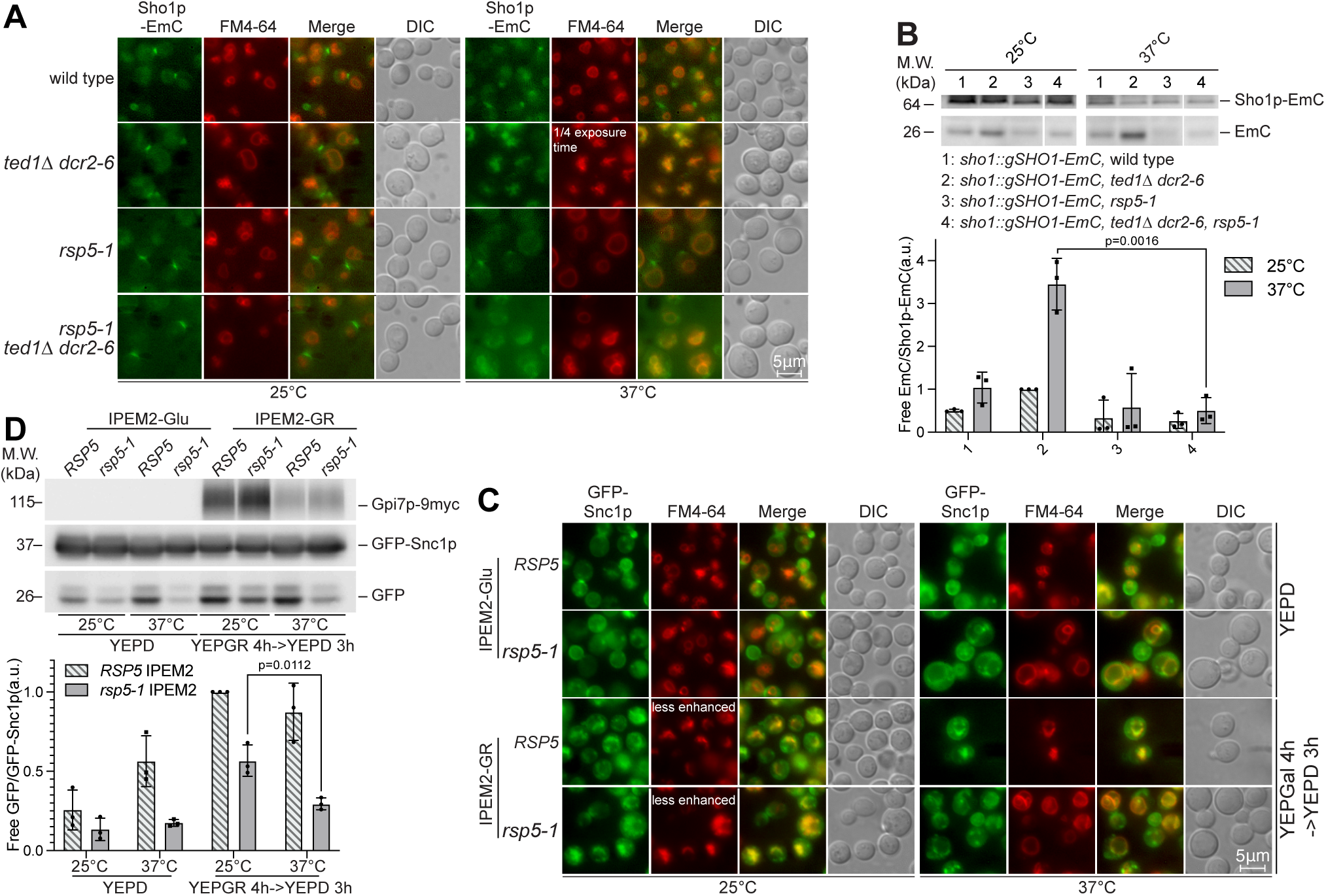
Rsp5p plays multiple roles in the endocytosis of proteins in GPI-AP Man2 remodeling mutants. (A) In *rsp5-1 ted111 dcr2-6* cells grown at 37°C Sho1p-EmC is reaches the limiting membrane of the vacuole. (B) In *rsp5-1 ted111 dcr2-6* cells grown at 37°C Sho1p-EmC is not internalized and degraded in the vacuole. WCEs from the indicated strains grown at 25 or 37°C were subjected to SDS-PAGE and immunoblotting. Pgk1p serves as a gel loading control. (C) The endocytosis and internalization of GFP-Snc1p in the vacuole is diminished in *rsp5-1* IPEM2-GR cells grown at 37°C. Note that the fragmented vacuole phenotype evident in IPEM2-GR cells is largely absent in *rsp5-1* IPEM2-GR cells grown at 37°C. (D) Immunoblots from WCEs of cells depicted in (C). Note that the degradation of GFP-Snc1p is largely blocked in *rsp5-1* IPEM2-GR cells grown at 37°C.

## Discussion

The plasma membrane is comprised of non-symmetrical distributions of various lipids and membrane proteins. However, the extent to which the formation of lipid domains on the plasma membrane is believed to involve the sequestration and subsequent enrichment of certain lipids (e.g., sphingolipids and sterols) has yet to be resolved. This uncertainty also applies to the role of and / or partitioning of GPI-APs within the plasma membrane (Simons and Sampaio, 2011; Simons and Toomre, 2000; Lingwood and Simons, 2007; Sevcsik et al., 2015). Nevertheless, the segregation of membrane proteins in plasma membrane is thought to play an important role in the creation of platforms that can mediate a variety of signalling events - activities that are critical to the cell’s capacity to respond to a variety of stimuli and as such vital for cellular adaptation and survival. (Lakhan and Sabharanjak, 2009; Levental et al., 2010; Suzuki et al., 2012; Sharonov et al., 2016; Szpurka et al., 2008; Zeng et al., 2023).

While it is well documented that removal of EtNP from Man2 functions as a transport warrant for the robust export of GPI-APs from the ER, there is currently limited information regarding the consequences of evading this remodeling event (Chen et al., 2021). In this study, we set out to further delineate the cell biological consequences of Man2 unremodeled GPI-APs trafficked to the plasma membrane of budding yeast cells. We show that in addition to activation of the SAC and the cell wall integrity pathway (Chen et al., 2021), unremodeled GPI-APs also trigger abnormal clathtrin-mediated endocytosis of certain membrane proteins and lead to the formation of numerous small vacuoles. Given the evolutionary ubiquity of the remodeling event, our findings may be broadly applicable to eukaryotic cells.

Despite the observation that some Man2 unremodeled GPI-APs, such as Gas1p and Sed1p, are not cleaved (Figure 2B and Figure S2B) and one of these, Gas1p, is not endocytosed (Figure 2C), we cannot exclude the possibly that other yeast GPI-APs are not endocytosed in IPEM2 cells (Yin et al., 2005). Indeed, others have shown that several yeast GPI-AP are subject to robust endocytosis and trafficking to the vacuole (MacDonald et al., 2015). As such, it seems most plausible that the phenotypes we have characterized here and elsewhere (Chen et al., 2021) for the IPEM2 mutant most likely arise from trafficking and delivery of GPI-APs bearing EtNP on Man2 to the plasma membrane. Consistent with this, we found that IPEM2 cells exhibit an increase in membrane disorder when grown under conditions were EtNP is not removed from Man2 of GPI-APs (Figure 1H and 1I).

Presumably, the accumulation of atypically larger numbers of GPI-APs in the plasma, such as in mutants where the cleavage the protein from the GPI is impaired, triggers a stress response. This response may be to disruption of signalling platforms, or to a perceived excess of GPI-APs outside of signalling platforms, which may impact lipid homeostasis (Figure 1H and 1I). In this regard, we note that the segregation of another so-called lipid raft protein (Pma1p) into DRMs was not affected in IPEM2-GR cells (Figure 1D), and that the signalling response pathway leading to activation of the cell wall integrity pathway also remained intact (Posas et al., 1998; Chen et al., 2021).

Yeast mutants defective in a mannosidase that transfers the GPI-linked protein to the cell wall reportedly exhibit a cell cycle phenotype, wherein cells arrest growth with small buds (Kitagaki et al., 2004). Similarly, *cdc1* mutants, which accumulate GPI-APs bearing EtNP on Man1, also arrest growth with small buds (Hartwell, 1971; Vazquez et al., 2014; Yang and Banfield, 2020). However, cells carrying mutations mannosidases that cleave GPI-APs (Figure S3G, S3H and S3I) do not phenocopy IPEM2 cells (Figure S3G, S3H and S3I). These data are in accord with our presumption that it is the failure to remodel GPI-APs, rather than defects in GPI-AP cleavage, that generates plasma membrane stress signals. Indeed, *cdc1* mutants display many of the phenotypes we have characterized for IPEM2 cells, which can perhaps best be reconciled with the view that Man1 unremodeled GPI-APs on the plasma membrane also elicit a stress response (Li, Lau, Yang and Banfield, unpublished observations).

Is the abnormal endocytosis observed in IPEM2-GR cells triggered in direct response to perturbations in the plasma membrane, or as part of an archetypal response whose purpose is to reset the organization and / or compositional status of the plasma membrane? Universal clearance of Man2 unremodeled GPI-APs from the plasma membrane may not be the objective of the stress-induced abnormal clathrin-mediated endocytosis we uncover here, as at least one GPI-AP, Gas1p, was not endocytosed (Figure 2C). Besides, others have shown some GPI-AP undergo robust endocytosis when properly remodeled (MacDonald et al., 2015). Furthermore, the fate of proteins that are endocytosed in IPEM2-GR cells (Sho1p and Snc1p) is distinct from that which occurs in otherwise wild type cells (Figure 3, Figure S3 and Figure 4). In wild type cells, Sho1p is rerouted from the plasma membrane to the bud neck, whereas Snc1p is recycled between the plasma membrane and the nascent bud. In IPEM2 cells, these proteins are endocytosed and redirected to the vacuole for degradation via the multivesicular body (Figure 3, Figure S3 and Figure 4). It seems plausible that the endocytic phenomena observed in IPEM2 mutant cells may have at least two purposes: to re-establish plasma membrane homeostasis and to transmit stress signals to the interior of the cell (Chen et al., 2021).

## Material and Methods

*S. cerevisiae* strains used in this study are listed in Table 1, plasmids used in this study are listed in Table 2 and antibodies used in this study are listed in the Reagents and Tools Table.

**Table 1.**
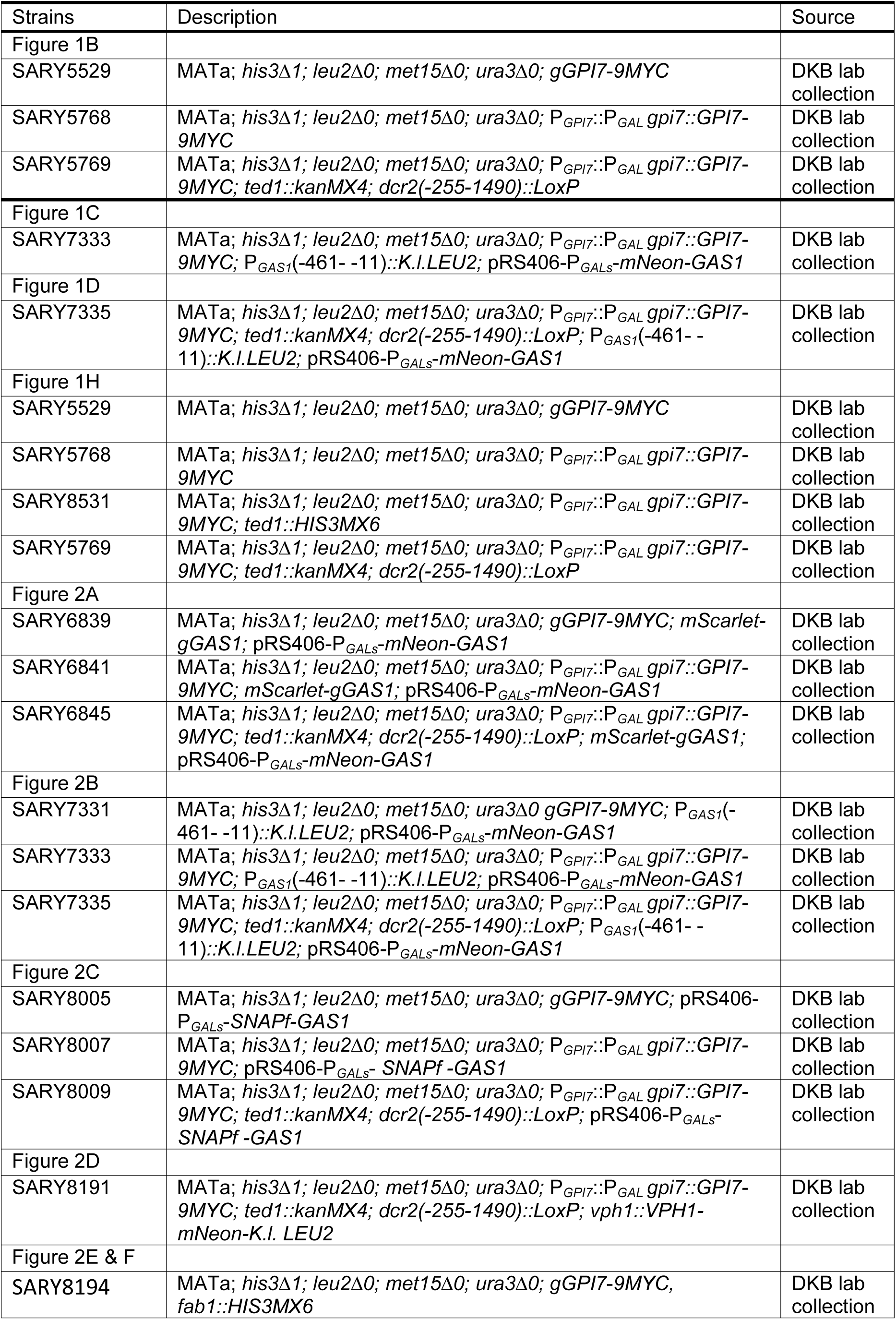
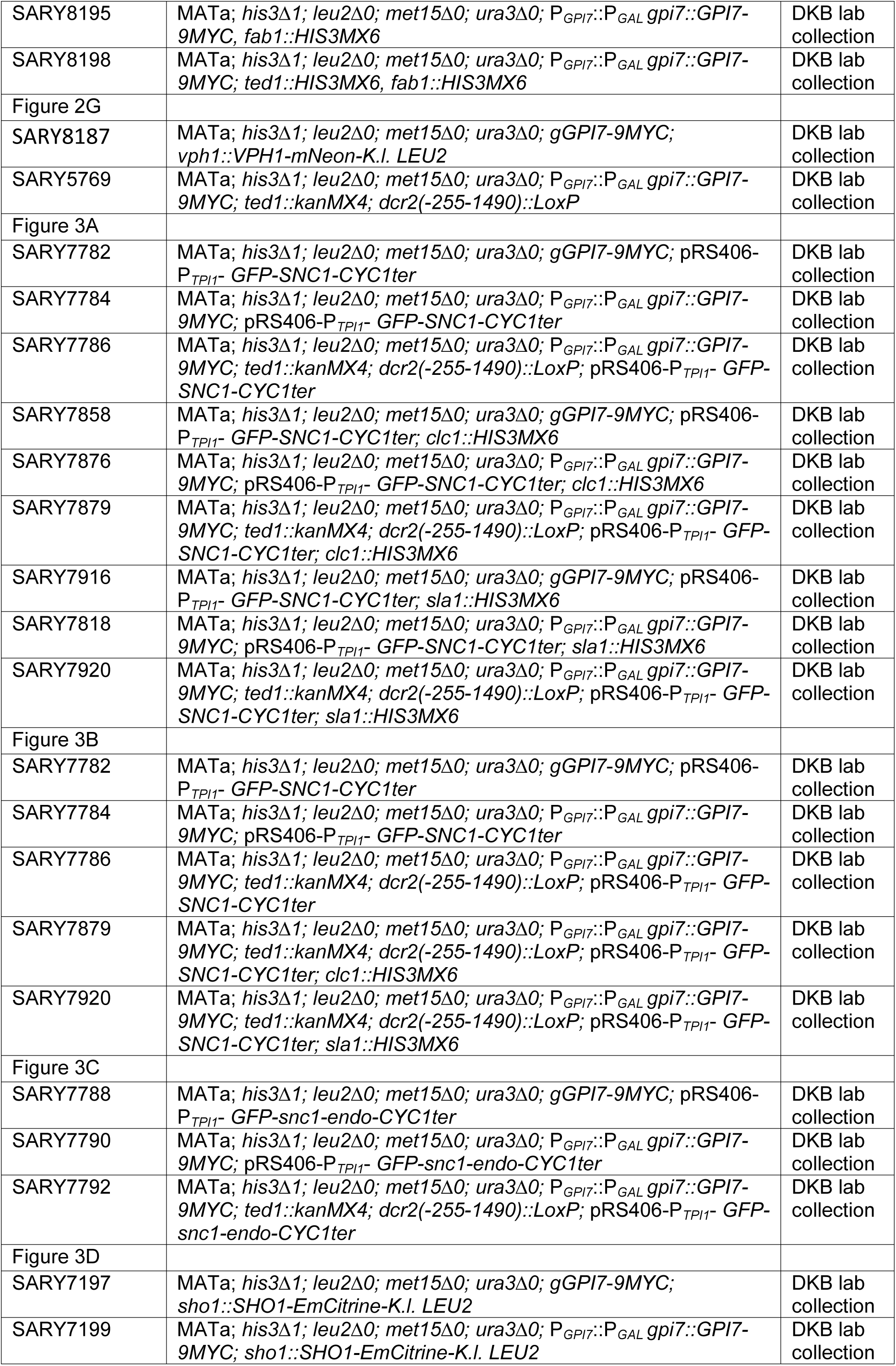
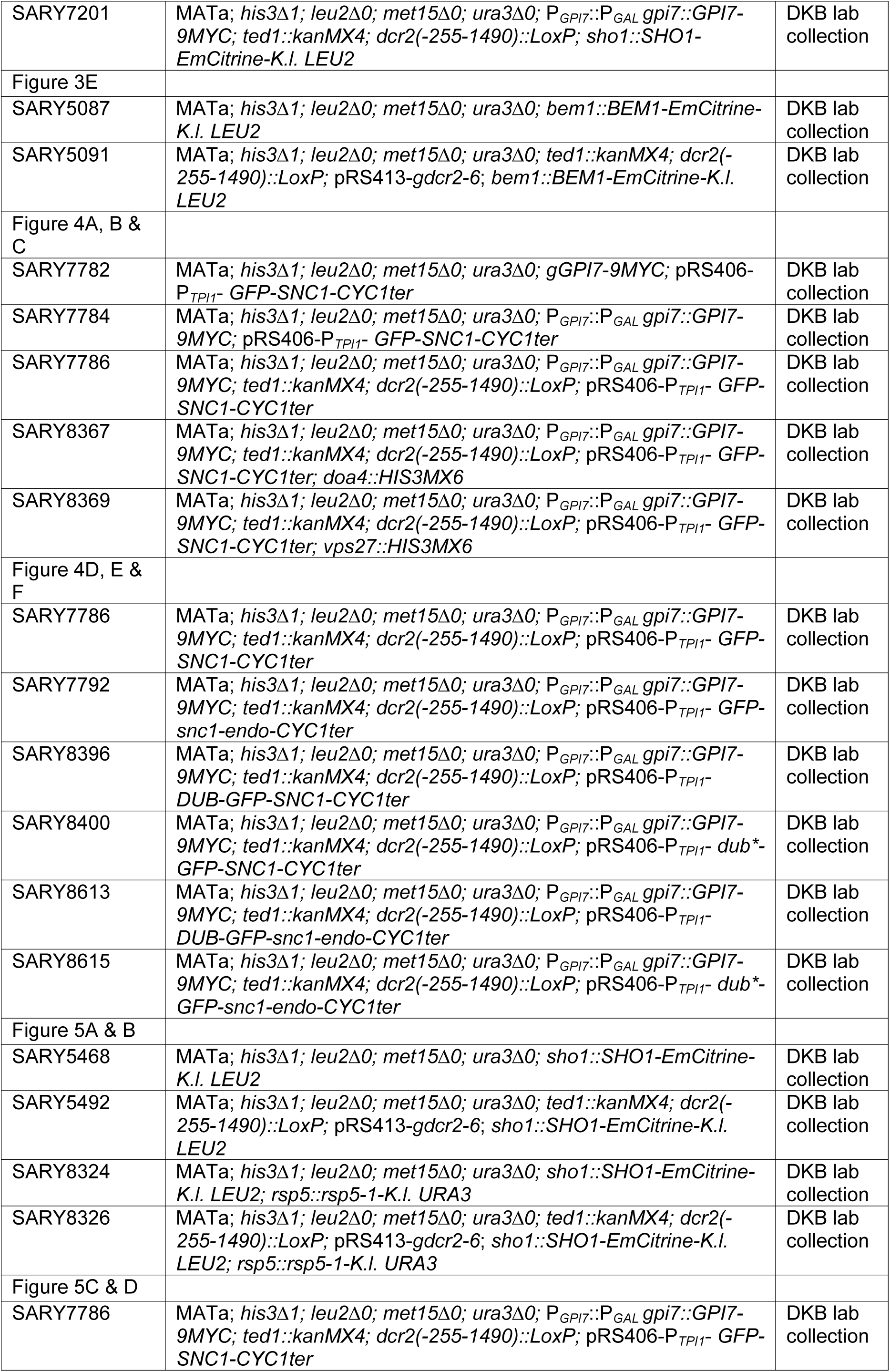
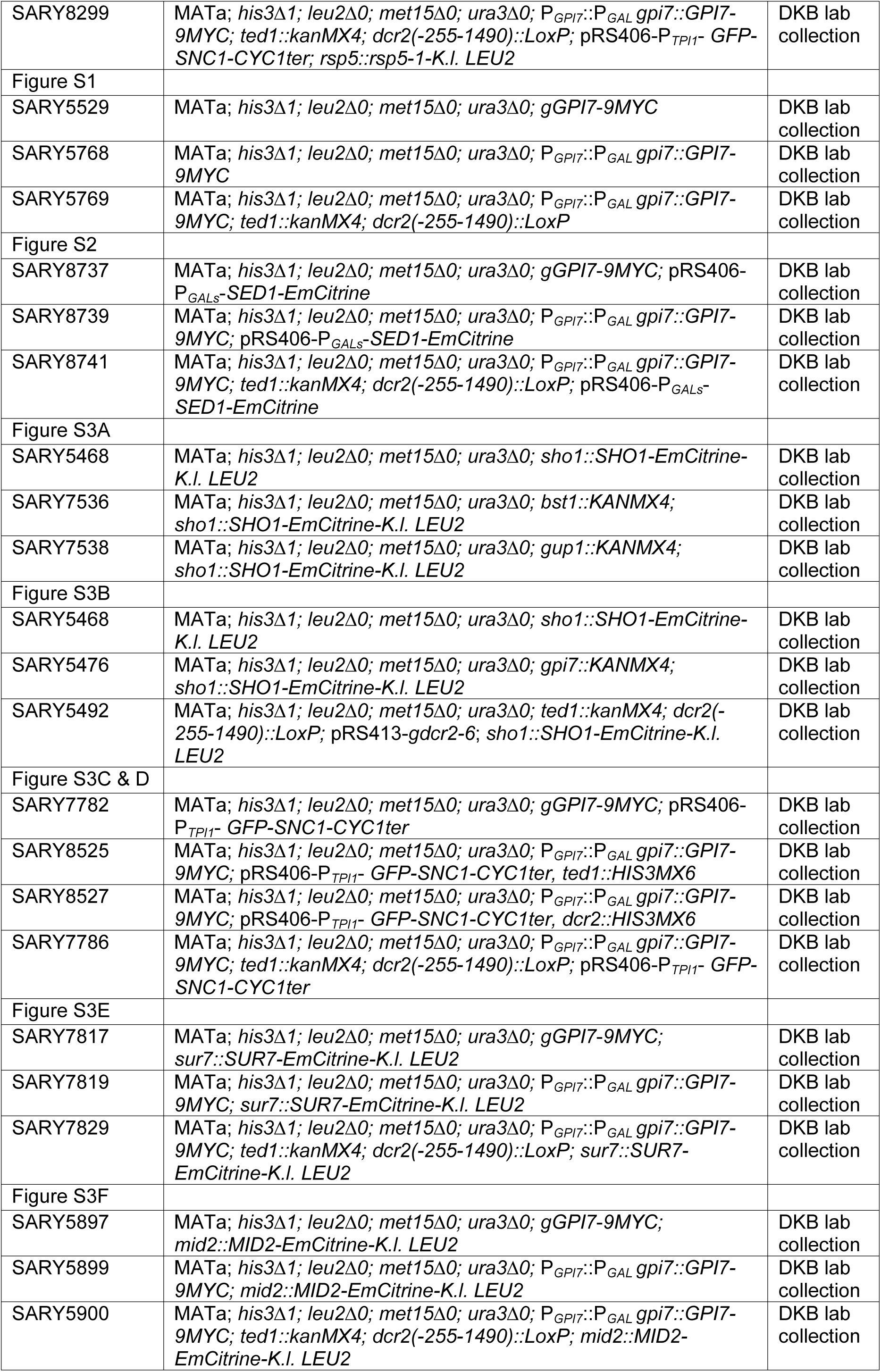
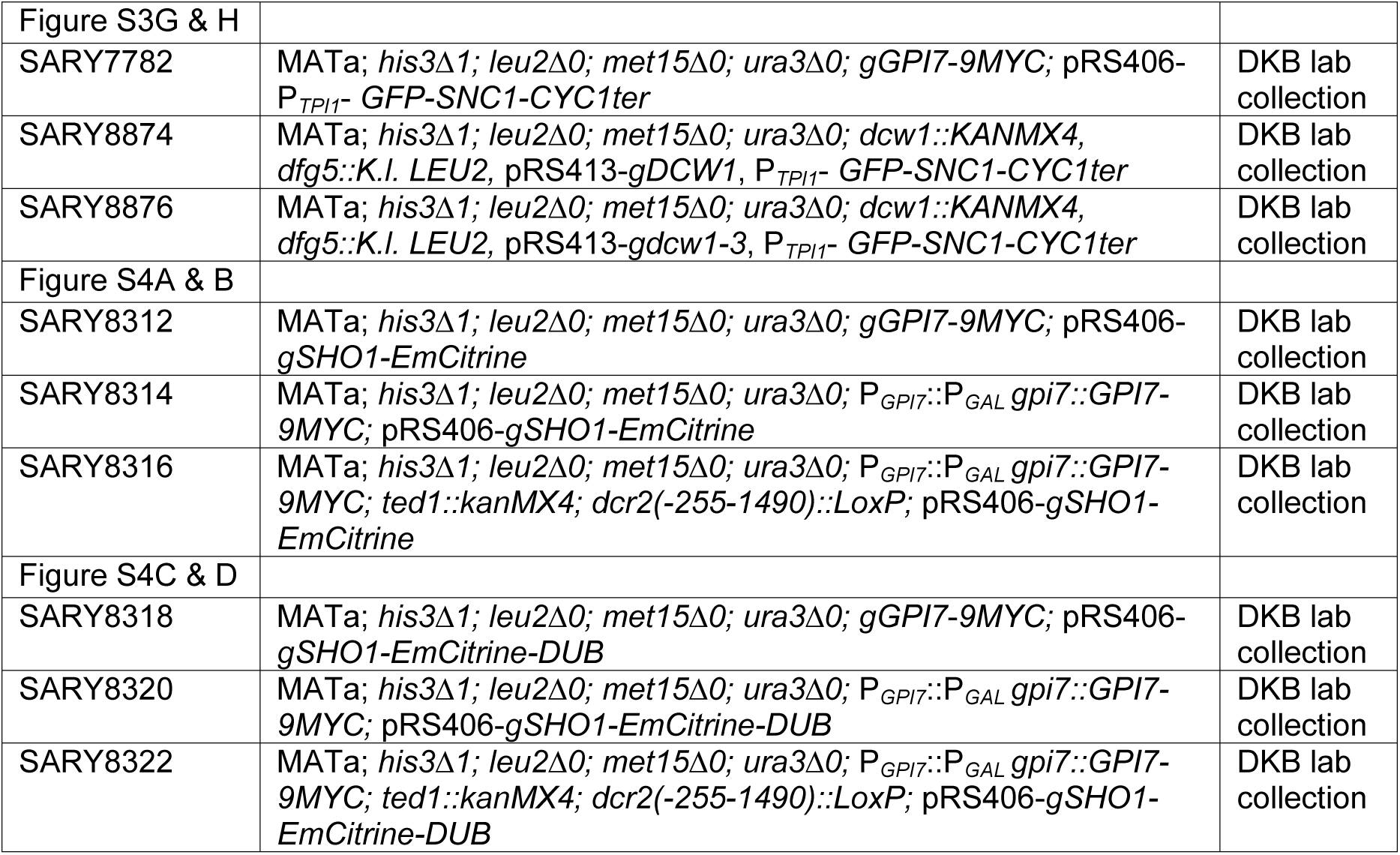
Strains used in this study.

**Table 2.**
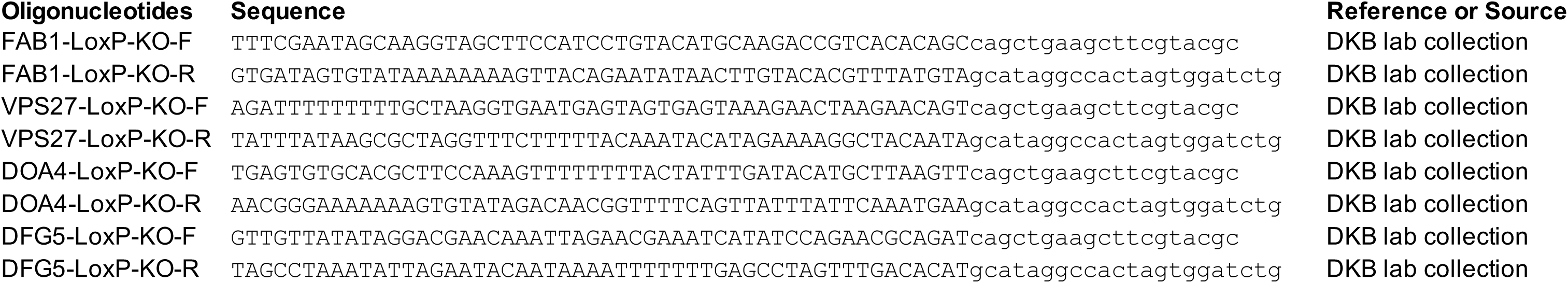
oligo used in this study.

## Experimental details

### Induction of Gpi7p synthesis

Induction of Gpi7p synthesis was conducted essentially as described in Chen and Banfield (2022). In brief, yeast strains were cultured in YEPD at 25°C overnight and thereafter diluted in YEPD to obtain a cell density of 0.3 X 10^7^ cells/mL and grown for a further three hours at 25°C. Cells were collected by centrifugation and washed once with YEP (i.e., in the absence of glucose), resuspended in YEPGR (containing 2% galactose (Sigma, Cat#G0750) and 1% raffinose (Sigma, Cat#R0250)) and incubated at 25°C for four hours to induce the expression of Gpi7p. Depending on the experimental design (see main text for details), cells were either cultured in YEPGR for an additional two hours (to achieve an ideal fluorescent signal intensity of GPI-AP markers), or were washed with YEP, resuspended in YEPD and incubated for an additional three hours at 25°C, to minimize the influence of Gpi7p over-expression and to recover the cells from a less preferable carbon source.

### Yeast cell viability test

1×10^7^ yeast cells were collected by centrifugation and then washed once with buffer (10mM HEPES, pH 7.2 containing 2% glucose). Cell pellets were resuspended in 1 mL of wash buffer to which 1 μL of Component A of from the LIVE/DEAD Yeast Viability Kit (Molecular Probe, Cat#L7009) and 5 μL of Component B was added. Cells suspended in the viability assay were incubated at 25°C in the dark for 30 minutes, washed once with wash buffer, and thereafter view by fluorescence microscopy (as per the manufacturer’s instructions). The percentage of viable cells was calculated as follows: percentage cell viability = (total # cells – total # dead cells) / total # cells x100.

### Detergent-Resistant Membrane (DRM) isolation

The DRM isolation procedure was performed based on the method described by Bagnat et al., (2000). In brief, 5×10^8^ cells were collected from log phase cultures and washed once with TNE buffer (50 mM Tris-HCl, pH7.4, 150 mM NaCl, 5 mM EDTA) before being stored at -80°C. Cells were thawed on ice and thereafter lysed at 4°C using acid washed glass beads (Sigma, Cat#G8772) in 1mL TNE buffer containing a protease inhibitor mixture 1mM Pefabloc SC (Roche, Cat#11429876001), 1×EDTA-free protease inhibitor cocktail (Roche, Cat#11873580001). Unbroken cells were removed by two rounds of centrifugation at 500xg for five minutes. The resulting lysate was adjusted to a final concentration of 1% Triton X-100 (Sigma, Cat#X100) and thereafter incubated on ice for 30 minutes. A total of 250 µL Triton X-100-treated lysate was mixed with 500 µL Optiprep Density Gradient Medium (Sigma, Cat#D1556) containing protease inhibitors generating a final 40% iodixanol mixture. Lysate (628 µL) was loaded onto the bottom of an ultracentrifugation tube (for a S55-S rotor, Thermo Scientific) and overlaid with 1005 µL of 50% Optiprep medium in TXNE buffer (TNE buffer containing 0.1% Triton X-100 and protease inhibitors). An additional 167 µL of TXNE buffer protease inhibitor mixture was layered onto the top of the sample bringing the final volume to 1800 µL. Samples were centrifuged at 55,000 rpm for 2 hours at 4°C in a S55-S rotor. Following centrifugation, six 300 µL fractions were collected from the top of the centrifugation tube. Proteins in each 300 µL fraction were precipitated by the addition of 600 µL ice cold 15% Trichloroacetic acid (TCA) (Sigma, Cat#T8657) and following mixing, each fraction was incubated at -20°C overnight. Precipitated proteins were collected by centrifugation in an Eppendorf microcentrifuge at 4°C and 16,100xg for 10 minutes. Sedimented proteins were washed once with cold acetone (-20°C) and thereafter dissolved in SDS-PAGE sample buffer, heated to 95°C for 5 minutes after which protein samples were subjected to SDS-PAGE and immunoblotting.

### Lipid order / disorder imaging and quantification

Lipid order / disorder imaging and quantification was performed essentially as described by Owen et al. (2011). In brief, a total of 10^7^ yeast cells were cultured in YEPD or in YEPGR for 4 hours and thereafter in YEPD for additional 3 hours. Yeast cells were collected by centrifugation, washed once with ice cold YEPD and then resuspended in 1mL ice cold YEPD containing 5 µM Di-4-ANEPPDHQ (Invitrogen, Cat#D36802). The cell Di-4-ANEPPDHQ suspensions were incubated on ice in the dark for 20 minutes, after which the stained cells were washed three times with ice cold PBS, mounted onto slides and viewed with a Leica SP8 confocal microscope.

A wavelength of 488 nm was used to excite Di-4-ANEPPDHQ and the emission signal was collected between 500-580 nm and 620-750 nm to acquire images corresponding to ordered and disordered lipids (respectively). The generalized polarization (GP) value was calculated using the following equation: GP=(I_500-580_-GI_620-750_)/ (I_500-580_+GI_620-750_). The G factor was calculated using the equation: G=(GP_ref_+GP_ref_GP_mes_-GP_mes_-1)/ (GP_mes_+GP_ref_GP_mes_-GP_ref_-1). The value of GP_ref_ was -0.85 for Di-4-ANEPPDHQ, while GP_mes_ was the GP value for a 5mM solution of Di-4-ANEPPDHQ dye in dimethyl sulfoxide (DMSO).

The GP value and the hue saturation brightness images were generated using the ImageJ macro provided by Owen, et al. (2011). Regions of interest were selected manually by adjusting the threshold of the gray scale image in ImageJ. The moving average trendline of the GP value histogram (Figure 1I) was added using Excel (Microsoft)

### Labelling vacuoles with FM4-64 dye

10^7^ yeast cells were cultured in YEPD or YEPGal for four hours, collected by centrifugation and resuspended in 50 μL of YEPD containing 30 μM FM 4-64 (Invitrogen, Cat#T13320). Yeast cells were incubated at 25°C for 30 minutes, harvested by centrifugation, and washed once with YEPD and thereafter incubated in 2 mL of YEPD at 25°C for 120 minutes before being subjected to epifluorescence microscopy. The Z-stack images of FM 4-64-labelled vacuoles from IPEM2 cell were obtained using confocal microscopy with a Zeiss LSM980.

### Monitoring endocytosis with FM 4-64 and SNAP-Surface® 488 dyes

Cultured yeast cells were harvested by centrifugation and resuspended in ice-cold PBS on ice for 10 minutes. Chilled cells were thereafter resuspended in ice-cold PBS containing 30 μM FM 4-64 (Invitrogen, Cat#T13320) and 5μM SNAP-Surface® 488 (NEB, Cat#S9124S), and incubated on ice for 30 minutes. Following incubation with FM 4-64 and SNAP-Surface® 488 cells were washed twice with ice-cold PBS to remove excess dyes. Yeast cells were either immediately mounted onto Concanavalin A (Sigma, Cat#C7275) coated slides for microscopy or cultured in YEPD medium for two hours (to monitor intracellular trafficking of FM 4-64 and SNAPf-Gas1p) before being subjected to confocal microscopy.

### Monitoring endocytosis of IPEM2-GR and Vph1p-mNeon-labelled wild type cells with FM 4-64 dye

IPEM2-GR and wild type cells were grown under identical conditions, collected by centrifugation. Equal numbers of cells from both strains were mixed together and resuspended in ice-cold PBS on ice for 10 minutes. Chilled cells were thereafter resuspended in ice-cold PBS containing 30 μM FM 4-64 (Invitrogen, Cat#T13320) and incubated on ice for a further 30 minutes. Following incubation with FM 4-64 cells were washed twice with ice-cold PBS to remove excess dyes. Yeast cells were immediately mounted onto a Concanavalin A (Sigma, Cat#C7275) coated 35mm confocal dish (SPL, Cat#100350) and covered with 2 mL of complete synthetic defined medium containing 2% glucose. Endocytosis was monitored, and images captured over the course of 160 mins by confocal microscopy with a Zeiss LSM980.

### Epifluorescence microscopy

Following yeast strain incubation cells were washed and resuspended in PBS. Aliquots of yeast cell suspensions (0.8 μL) were placed onto slides coated with Concanavalin A (Sigma, Cat#7275) and examined by epifluorescence microscopy. Cells were photographed immediately after examination.

Cell visualization and photography were performed using a Nikon ECLIPSE 80i microscope (Nikon Instruments, Japan) equipped with a Nikon Plan Apo VC 100X/1.40 oil objective lens and a SPOT-RT3 monochrome camera (DIAGNOSTIC Instruments, Inc., Sterling Heights, MI). Images were acquired using SPOTSOFTWARE (version 4.6, Diagnostic Instrument, Inc). Digital images were processed using Photoshop CS6 software (Adobe Systems, Mountain View, CA).

### Sample preparation for immunoblotting

10^7^ yeast cells were collected by centrifugation and thereafter resuspended in 15% Trichloroacetic acid (TCA) (Sigma, Cat#T8657). The cell – TCA suspension was stored at -20°C overnight after which the resulting precipitate was collected by centrifugation and washed once with cold acetone. After being air-dried, the precipitate was dissolved in SDS-PAGE sample buffer containing 50 mM NaOH and 0.5% SDS. The sample was then subjected to SDS-PAGE and transferred to a nitrocellulose membrane overnight using a current of 250 mA.

### Quantification of immunoblots

For Figure 1E, the ratio (mGas1p/pGas1p) from each fraction in Figures 1C and 1D was calculated using the formula: Ratio = I_m_/I_p_. Here, “I_m_” represents the signal intensity of mNeon-mGas1p in each fraction, and “I_p_” represents the signal intensity of mNeon-pGas1p in each fraction.

The percentage of mNeon-mGas1p from each fraction in Figures 1C and 1D was calculated using the formula: Percentage = I_n_/I_total_ (x100). In this formula, “I_n_” represents the signal intensity of mNeon-mGas1p in each fraction, while “I_total_” is the sum of the signal intensity of mNeon-mGas1p from all six fractions (Figure 1F).

To calculate the ratio (s-ALP/m-ALP), the following formula was used: Ratio = I_s_/I_m_. Here, “I_s_” represents the signal intensity of s-ALP in each fraction (fractions 4-6), and “I_m_” represents the signal intensity of m-ALP in each fraction (fractions 4-6) (Figure 1G).

The ratio (in arbitrary units, a.u.) of free GFP (Figures 3B, 4C, 4E and 4F) was calculated using the following formula: Ratio = I_GFP_/I_GFP-Snc1p_. In these formulas, “I_GFP_” refers to the signal intensity of free GFP in each sample, “I_GFP-Snc1p_” refers to the signal intensity of GFP-Snc1p in the same sample.

### Quantification of protein colocalization

The calculation of Pearson’s coefficient was performed using the JACoP ImageJ plugin (Bolte and Cordelieres, 2006). The regions of cells of interest were cropped and subjected to JACoP, and Costes’ automatic threshold was applied to the paired cropped images to calculate the Pearson’s coefficient.

## Acknowledgements

We thank the Biosciences Central Research Facility (BioCRF) at HKUST (Clear Water Bay) for the use of the Leica SP8 confocal microscope and the ChemiDoc system. This work was supported in part by grants from the Hong Kong Research Grants Council to DK Banfield (16102320 and 16102722)

## Author information

Authors and Affiliations

**The Division of Life Science, The Hong Kong University of Science and Technology. Clear Water Bay, Kowloon, Hong Kong, Special Administrative Region of China**

Li Chen and David K. Banfield

## Contributions

L. Chen and DK Banfield conceived project, DK Banfield supervised project, L. Chen performed experiments, L. Chen and DK Banfield analysed data, DK Banfield wrote the manuscript with input from L. Chen.

## Corresponding author

Correspondence to DK Banfield

## Supplementary Figure Legends

**Figure S1.**
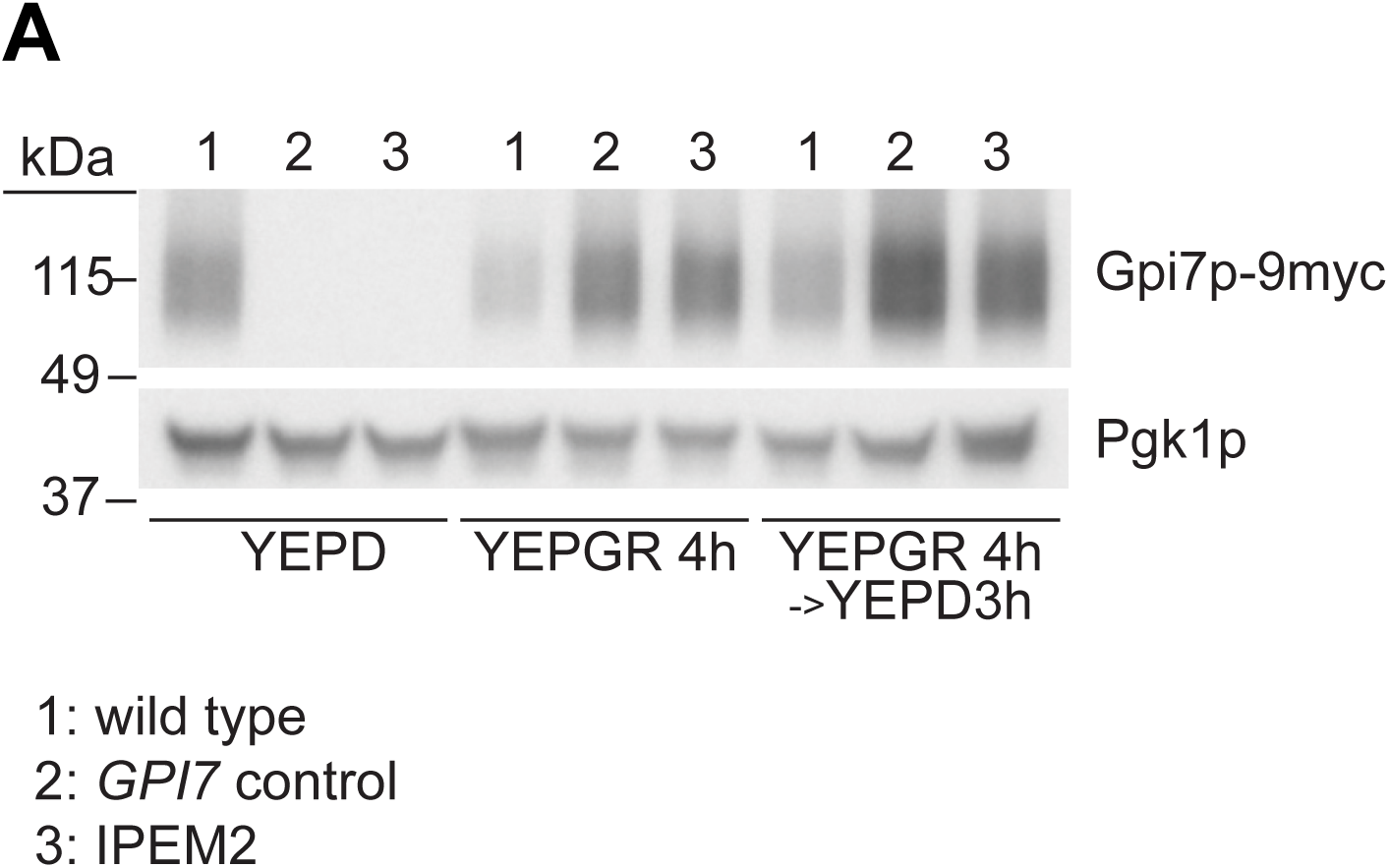
The expression profile of *GPI7* in IPEM2-GR cells and control strains. *GPI7* was expressed as a C-terminal fusion protein to 9 copies of the myc epitope using either the *GPI7* promoter or the *GAL 1/10* promoter, as indicated. WCEs from the indicated strains grown under the conditions indicated were subjected to SDS-PAGE and immunoblotting. Pgk1p serves as a gel loading control.

**Figure S2.**
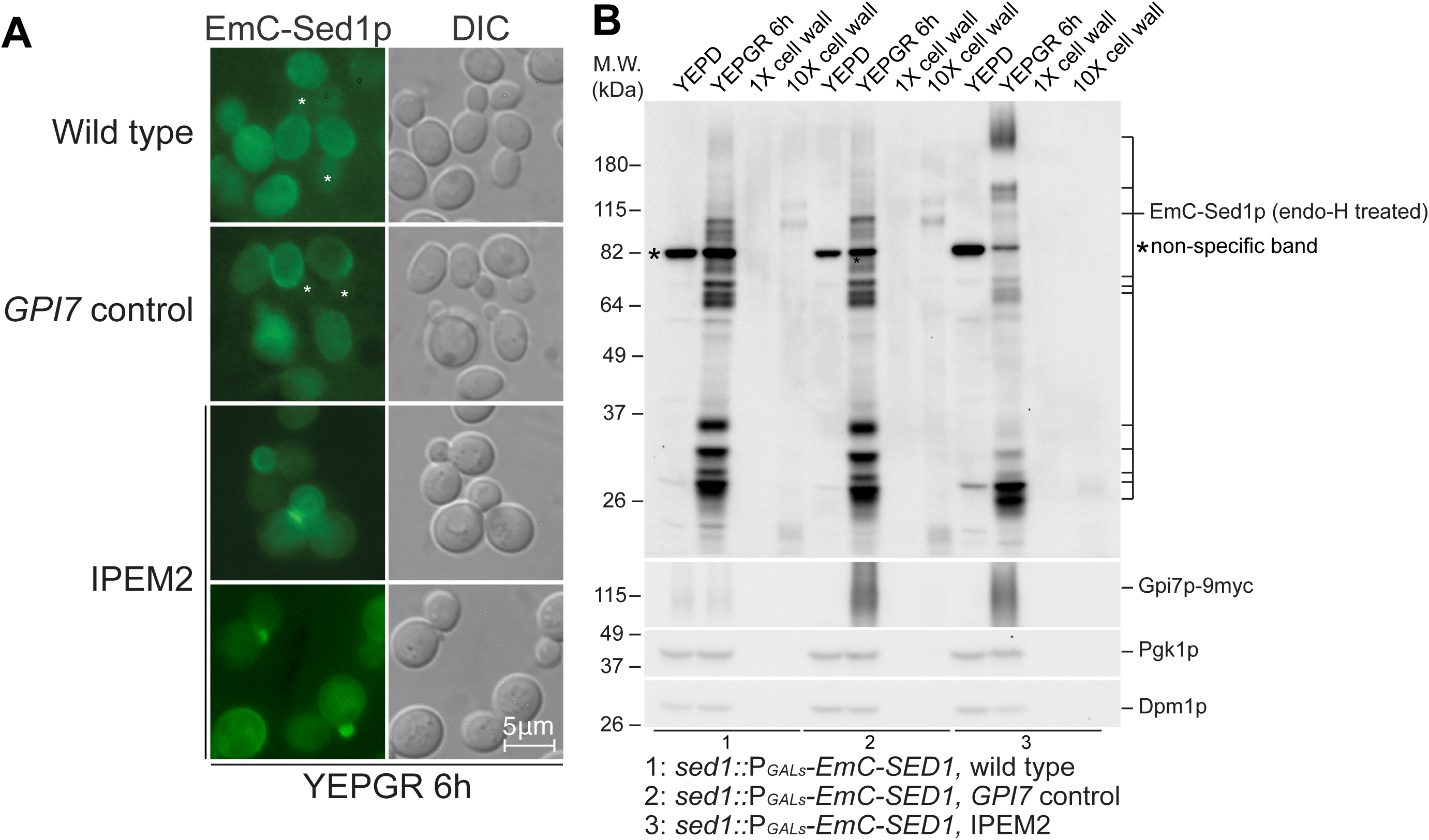
In IPEM2-GR cells Sed1p is not cleaved and crosslinked to the cell wall. (A) The distribution of EmC-Sed1p in IPEM2-GR cells and control strains. EmC (enhanced monomeric citrine). Note the exclusion of *de novo* synthesized EmC-Sed1p from the daughter cell in wild type and control cells (asterisk), and the presence of newly synthesized EmC-Sed1p in the nascent buds of IPEM2-GR cells. (B) In IPEM2-GR cells EmC-Sed1p is not associated with the cell wall. EmC-Sed1p was detected using an anti-GFP antibody. *GPI7* expression was monitored with an anti-myc antibody, whilst the robustness of the separation of the cell wall from cellular components was assessed by immunostaining with antibodies against a cytosolic protein (Pgk1p) and an ER resident protein (Dpm1p).

**Figure S3.**
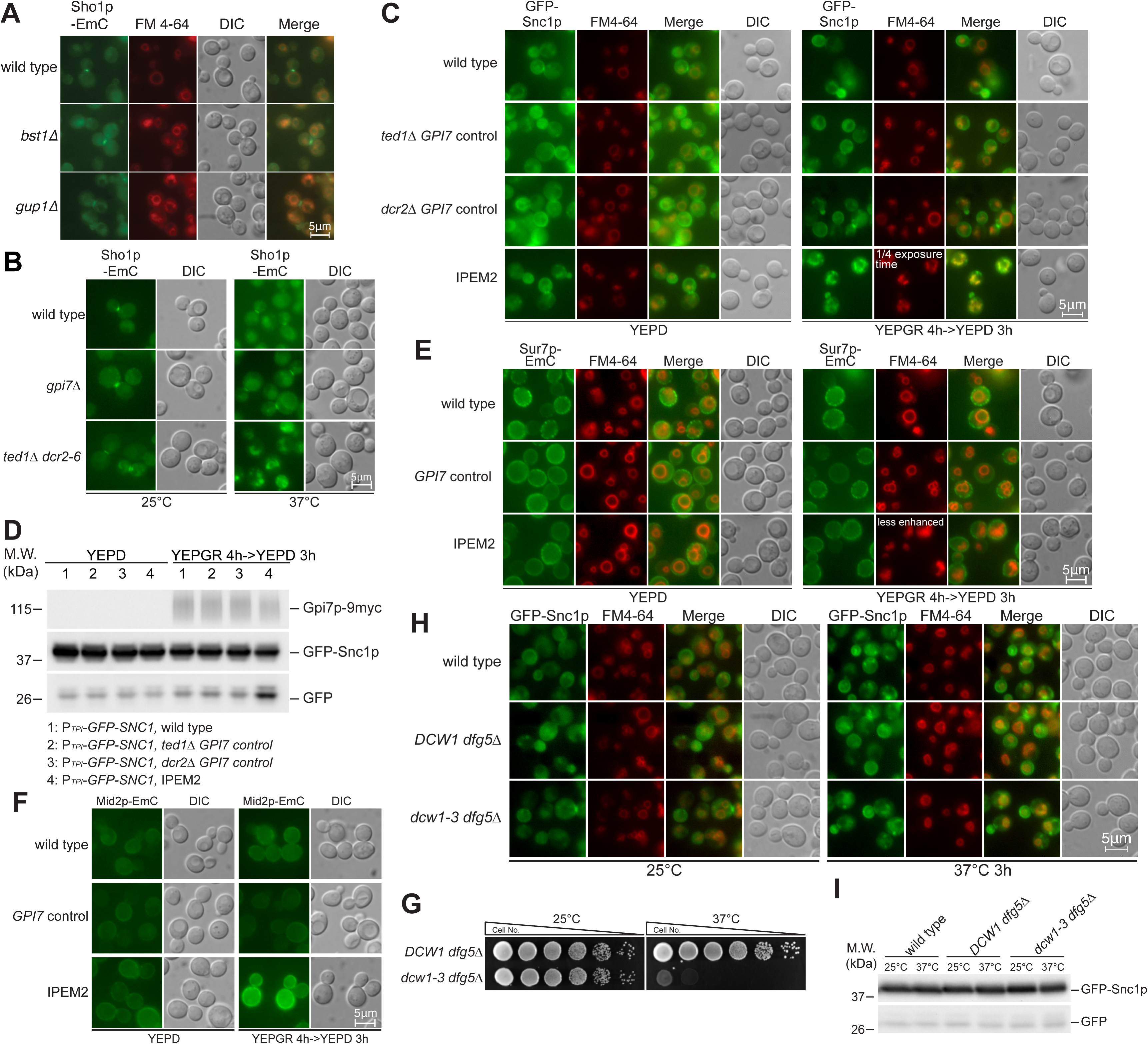
Endocytic trafficking of Sho1p, Snc1p, Sur7p and Mid2p in GPI-AP remodeling mutants and in IPEM2-GR cells. (A) Sho1p-EmC is not mislocalized to the vacuole in cells lacking the GPI-AP remodelase genes *BST1* and *GUP1*. (B) Sho1p-EmC is largely restricted to the bud neck and limiting membrane of the vacuole in cells lacking *GPI7* grown at 37°C but localizes to numerous small vacuoles in IPEM2-GR cells. (C) The trafficking repertoire of GFP-Snc1p in cells lacking either *TED1* or *DCR2* is indistinguishable from that in wild type cells (D) GFP-Snc1p is largely excluded from the lumen of the vacuole in cells lacking either *TED1* or *DCR2*. (E) The intracellular distribution of the eisosome component Sur7p-EmC is unaffected in IPEM2-GR cells. (F) The intracellular distribution of the cell wall integrity pathway sensor Mid2p-EmC is unaffected in IPEM2-GR cells. Note that, as reported previously (Chen et al., 2021) the expression level of Mid2p-EmC is increased in IPEM2-GR cells. (G) The trafficking repertoire of GFP-Snc1p is unaffected in *dcw1-3 dfg511* cells grown at 37°C. (H) *dcw1-3 dfg511* cells are temperature-sensitive for growth. 10-fold serial dilutions of the indicated strains were spotted onto YEPD plates which were thereafter incubated at 25 or 37°C for 3 days prior to being photographed. (I) GFP-Snc1p is not degraded in *dcw1-3 dfg511* cells grown at 37°C.

**Figure S4.**
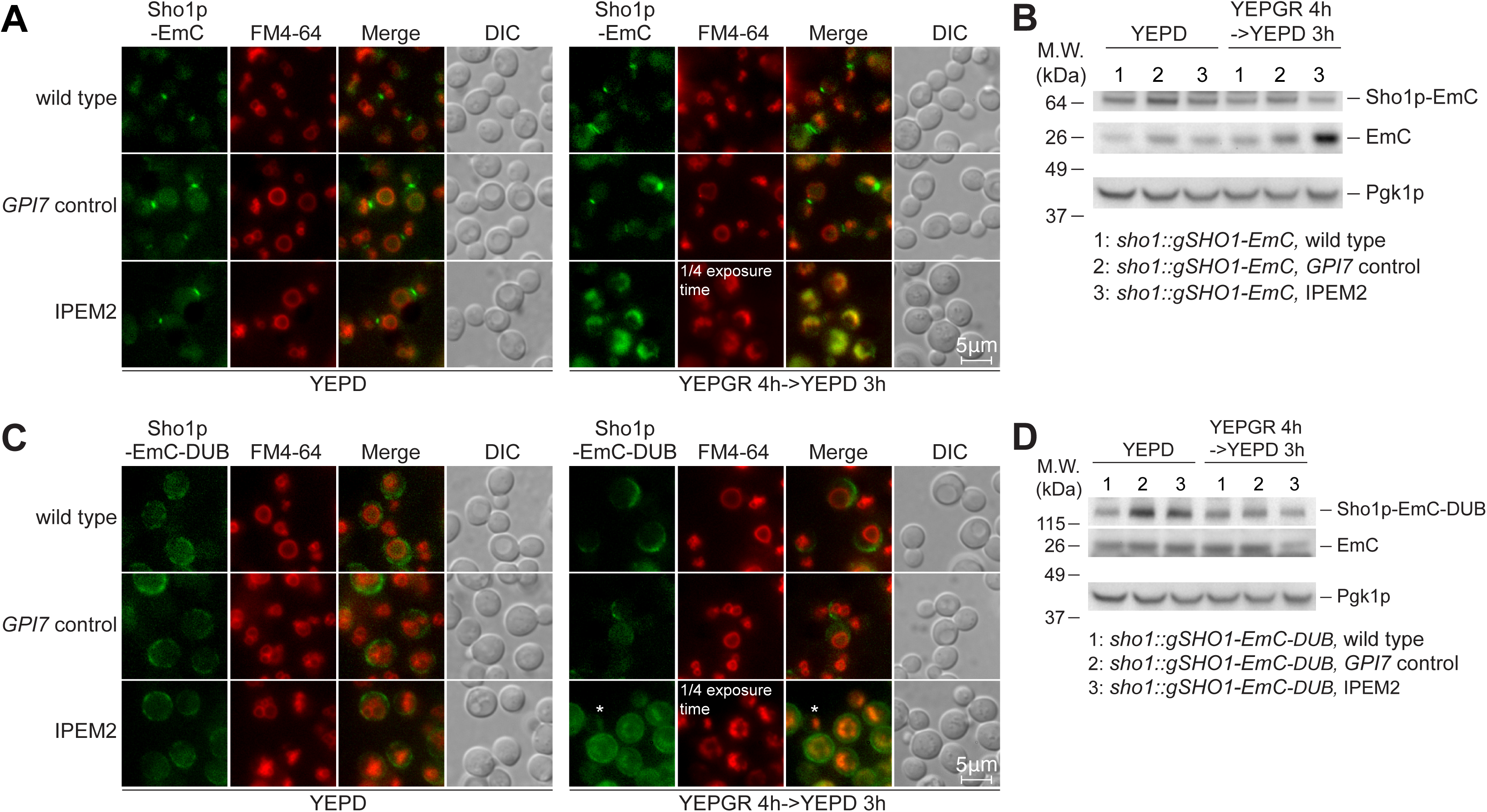
Ubiquitination of Sho1p-EmC is a prerequisite for the protein’s endocytosis in IPEM2-GR cells. (A) and (B) Sho1p-EmC is mislocalized to and degraded in numerous small vacuoles in IPEM2-GR cells. (C) and (D) The de-ubiquitin domain Sho1p-EmC fusion protein Sho1p-EmC-DUB is largely localized to the plasma membrane in IPEM2-GR cells. In panels (B) and (C) the cytoplasmic protein Pgk1p serves as a gel loading control.

## Structured Methods

### Reagents and Tools Table

Structured Methods Reagents and Tools Table
| Reagent/Resource | Reference or Source | Identifier or Catalog Number |
| --- | --- | --- |
| **Experimental Models** |  |  |
| *Saccharomyces cerevisiae* (BY4741) | DKB lab collection | N/A |
| Strains used in this study | DKB lab collection | Table 1 |
| **Recombinant DNA** |  |  |
| pRS406-P<sub>GALS</sub>-mNeon-GAS1 | DKB lab collection | N/A |
| pRS406-P<sub>GALS</sub>-EmCitrine-SED1 | DKB lab collection | N/A |
| pRS406-P<sub>GALS</sub>-SNAPf-GAS1 | DKB lab collection | N/A |
| pRS406-P<sub>TPI1</sub>-GFP-SNC1-CYC1ter | DKB lab collection | N/A |
| pRS406-P<sub>TPI1</sub>-GFP-snc1-endo-CYC1ter | DKB lab collection | N/A |
| pRS406-P<sub>TPI1</sub>-DUB-GFP-SNC1-CYC1ter | DKB lab collection | N/A |
| pRS406-P<sub>TPI1</sub>-dub*-GFP-SNC1-CYC1ter | DKB lab collection | N/A |
| **Antibodies** |  |  |
| Rabbit polyclonal anti Gas1p | kindly provided by Prof. Howard Reizman (University of Geneva) | N/A |
| Mouse monoclonal anti ALP | Invitrogen | Cat# A6458 |
| Mouse monoclonal anti Pma1p | Invitrogen | Cat# MA1-91567 |
| Mouse monoclonal anti Pgk1p | Molecular Probes | Cat#459250 |
| Rabbit polyclonal anti GFP | Sigma | Cat#G1544 |
| Mouse monoclonal anti Myc | Roche | Cat#11667149001 |
| Mouse monoclonal anti Dpm1 | Invitrogen | Cat#A-6429 |
| Donkey Anti-Rabbit IgG | GE Healthcare | Cat#NA934 |
| Goat Anti-Mouse IgG | Sigma | Cat#A8924 |
| **Oligonucleotides and other sequence-based reagents** |  |  |
| PCR primers | This study | Table 2 |
| **Chemicals, Enzymes and** |  |  |
| **other reagents** |  |  |
| galactose | Sigma | Cat # G0750 |
| raffinose | Sigma | Cat # R0250 |
| LIVE/DEAD Yeast Viability Kit | Molecular Probe | Cat # L7009 |
| acid washed glass beads | Sigma | Cat # G8772 |
| Pefabloc SC | Roche | Cat # 11429876001 |
| EDTA-free protease inhibitor cocktail | Roche | Cat # 11873580001 |
| Triton X-100 | Sigma | Cat # X100 |
| Optiprep Density Gradient Medium | Sigma | Cat # D1556 |
| Trichloroacetic acid | Sigma | Cat # T8657 |
| Di-4-ANEPPDHQ | Invitrogen | Cat # D36802 |
| FM 4-64 | Invitrogen | Cat # T13320 |
| SNAP-Surface® 488 | New England Biolabs | Cat # S9124S |
| Concanavalin A | Sigma | Cat # C7275 |
| **Software** |  |  |
| Fiji | https://fiji.sc |  |
| Adobe Photpshop 2024 | https://www.adobe.com |  |
| Adobe Illustrator 2024 | https://www.adobe.com |  |
| Prism 10 | https://www.graphpad.com |  |
| Microsoft Office 365 | https://www.microsoft.com |  |
| **Other** |  |  |
| LSM980 | Zeiss |  |

## References

Amaro, M, Reina, F, Hof, M, Eggeling, C, and Sezgin, E (2017). Laurdan and Di-4-ANEPPDHQ probe different properties of the membrane. J Phys D Appl Phys 50, 134004.

Babst, M (2011). MVB vesicle formation: ESCRT-dependent, ESCRT-independent and everything in between. Curr Opin Cell Biol 23, 452–457.

Bagnat, M, Chang, A, and Simons, K (2001). Plasma membrane proton ATPase Pma1p requires raft association for surface delivery in yeast. Mol Biol Cell 12, 4129–4138.

Bagnat, M, Keränen, S, Shevchenko, A, Shevchenko, A, and Simons, K (2000). Lipid rafts function in biosynthetic delivery of proteins to the cell surface in yeast. Proc Natl Acad Sci U S A 97, 3254–3259.

Benachour, A, Sipos, G, Flury, I, Reggiori, F, Canivenc-Gansel, E, Vionnet, C, Conzelmann, A, and Benghezal, M (1999). Deletion of GPI7, a yeast gene required for addition of a side chain to the glycosylphosphatidylinositol (GPI) core structure, affects GPI protein transport, remodeling, and cell wall integrity. J Biol Chem 274, 15251–15261.

Bolte, S, and Cordelières, FP (2006a). A guided tour into subcellular colocalization analysis in light microscopy. J Microsc 224, 213–232.

Bolte, S, and Cordelières, FP (2006b). A guided tour into subcellular colocalization analysis in light microscopy. J Microsc 224, 213–232.

Bosson, R, Jaquenoud, M, and Conzelmann, A (2006). GUP1 of Saccharomyces cerevisiae encodes an O-acyltransferase involved in remodeling of the GPI anchor. Mol Biol Cell 17, 2636– 2645.

Chen, L, and Banfield, DK (2022). Engineering yeast to induce the synthesis of GPI-APs with a permanent phosphoethanolamine on mannose 2 of the glycan moiety. STAR Protoc 3, 101503.

Chen, L, Tu, L, Yang, G, and Banfield, DK (2021). Remodeling-defective GPI-anchored proteins on the plasma membrane activate the spindle assembly checkpoint. Cell Reports 37, 110120.

Dunn, R, and Hicke, L (2001). Domains of the Rsp5 Ubiquitin-Protein Ligase Required for Receptor-mediated and Fluid-Phase Endocytosis. MBoC 12, 421–435.

Fujihara, Y, and Ikawa, M (2016). GPI-AP release in cellular, developmental, and reproductive biology. J Lipid Res 57, 538–545.

Fujita, M, Maeda, Y, Ra, M, Yamaguchi, Y, Taguchi, R, and Kinoshita, T (2009). GPI glycan remodeling by PGAP5 regulates transport of GPI-anchored proteins from the ER to the Golgi. Cell 139, 352–365.

Fujita, M, Yoko-O, T, and Jigami, Y (2006). Inositol deacylation by Bst1p is required for the quality control of glycosylphosphatidylinositol-anchored proteins. Mol Biol Cell 17, 834–850.

Galluzzi, L, Yamazaki, T, and Kroemer, G (2018). Linking cellular stress responses to systemic homeostasis. Nat Rev Mol Cell Biol 19, 731–745.

Hartwell, LH (1971). Genetic control of the cell division cycle in yeast. IV. Genes controlling bud emergence and cytokinesis. Exp Cell Res 69, 265–276.

Hurst, LR, and Fratti, RA (2020). Lipid Rafts, Sphingolipids, and Ergosterol in Yeast Vacuole Fusion and Maturation. Front Cell Dev Biol 8, 539.

Jin, L, Millard, AC, Wuskell, JP, Clark, HA, and Loew, LM (2005). Cholesterol-enriched lipid domains can be visualized by di-4-ANEPPDHQ with linear and nonlinear optics. Biophys J 89, L04–06.

Katzmann, DJ, Sarkar, S, Chu, T, Audhya, A, and Emr, SD (2004). Multivesicular body sorting: ubiquitin ligase Rsp5 is required for the modification and sorting of carboxypeptidase S. Mol Biol Cell 15, 468–480.

Katzmann, DJ, Stefan, CJ, Babst, M, and Emr, SD (2003). Vps27 recruits ESCRT machinery to endosomes during MVB sorting. J Cell Biol 162, 413–423.

Keppler, A, Pick, H, Arrivoli, C, Vogel, H, and Johnsson, K (2004). Labeling of fusion proteins with synthetic fluorophores in live cells. Proc Natl Acad Sci U S A 101, 9955–9959.

Kinoshita, T (2020). Biosynthesis and biology of mammalian GPI-anchored proteins. Open Biol 10, 190290.

Kinoshita, T, and Fujita, M (2016). Biosynthesis of GPI-anchored proteins: special emphasis on GPI lipid remodeling. J Lipid Res 57, 6–24.

Kitagaki, H, Ito, K, and Shimoi, H (2004). A temperature-sensitive dcw1 mutant of Saccharomyces cerevisiae is cell cycle arrested with small buds which have aberrant cell walls. Eukaryot Cell 3, 1297–1306.

Kitagaki, H, Wu, H, Shimoi, H, and Ito, K (2002). Two homologous genes, DCW1 (YKL046c) and DFG5, are essential for cell growth and encode glycosylphosphatidylinositol (GPI)-anchored membrane proteins required for cell wall biogenesis in Saccharomyces cerevisiae. Mol Microbiol 46, 1011–1022.

Klionsky, DJ, and Emr, SD (1989). Membrane protein sorting: biosynthesis, transport and processing of yeast vacuolar alkaline phosphatase. EMBO J 8, 2241–2250.

Lakhan, SE, Sabharanjak, S, and De, A (2009). Endocytosis of glycosylphosphatidylinositol-anchored proteins. J Biomed Sci 16, 93.

Laude, AJ, and Prior, IA (2004). Plasma membrane microdomains: organization, function and trafficking. Mol Membr Biol 21, 193–205.

Leeuw, T, Fourest-Lieuvin, A, Wu, C, Chenevert, J, Clark, K, Whiteway, M, Thomas, DY, and Leberer, E (1995). Pheromone response in yeast: association of Bem1p with proteins of the MAP kinase cascade and actin. Science 270, 1210–1213.

Levental, I, Grzybek, M, and Simons, K (2010). Greasing their way: lipid modifications determine protein association with membrane rafts. Biochemistry 49, 6305–6316.

Lingwood, D, and Simons, K (2007). Detergent resistance as a tool in membrane research. Nat Protoc 2, 2159–2165.

MacDonald, C, Payne, JA, Aboian, M, Smith, W, Katzmann, DJ, and Piper, RC (2015). A Family of Tetraspans Organizes Cargo for Sorting into Multivesicular Bodies. Developmental Cell 33, 328–342.

Manzano-Lopez, J, Perez-Linero, AM, Aguilera-Romero, A, Martin, ME, Okano, T, Silva, DV, Seeberger, PH, Riezman, H, Funato, K, Goder, V, et al. (2015). COPII coat composition is actively regulated by luminal cargo maturation. Curr Biol 25, 152–162.

Michaillat, L, and Mayer, A (2013). Identification of Genes Affecting Vacuole Membrane Fragmentation in Saccharomyces cerevisiae. PLOS ONE 8, e54160.

Muñiz, M, and Zurzolo, C (2014). Sorting of GPI-anchored proteins from yeast to mammals--common pathways at different sites? J Cell Sci 127, 2793–2801.

de Nadal, E, and Posas, F (2022). The HOG pathway and the regulation of osmoadaptive responses in yeast. FEMS Yeast Res 22, foac013.

Nikko, E, and André, B (2007). Evidence for a direct role of the Doa4 deubiquitinating enzyme in protein sorting into the MVB pathway. Traffic 8, 566–581.

Nuoffer, C, Jenö, P, Conzelmann, A, and Riezman, H (1991). Determinants for glycophospholipid anchoring of the Saccharomyces cerevisiae GAS1 protein to the plasma membrane. Mol Cell Biol 11, 27–37.

Owen, DM, Rentero, C, Magenau, A, Abu-Siniyeh, A, and Gaus, K (2011). Quantitative imaging of membrane lipid order in cells and organisms. Nat Protoc 7, 24–35.

Paladino, S, Lebreton, S, and Zurzolo, C (2015). Trafficking and Membrane Organization of GPI-Anchored Proteins in Health and Diseases. Curr Top Membr 75, 269–303.

Parton, RG, and Hancock, JF (2004). Lipid rafts and plasma membrane microorganization: insights from Ras. Trends Cell Biol 14, 141–147.

Posas, F, Takekawa, M, and Saito, H (1998). Signal transduction by MAP kinase cascades in budding yeast. Curr Opin Microbiol 1, 175–182.

Satpute-Krishnan, P, Ajinkya, M, Bhat, S, Itakura, E, Hegde, RS, and Lippincott-Schwartz, J (2014). ER stress-induced clearance of misfolded GPI-anchored proteins via the secretory pathway. Cell 158, 522–533.

Sevcsik, E, Brameshuber, M, Fölser, M, Weghuber, J, Honigmann, A, and Schütz, GJ (2015). GPI-anchored proteins do not reside in ordered domains in the live cell plasma membrane. Nat Commun 6, 6969.

Sezgin, E, Levental, I, Mayor, S, and Eggeling, C (2017). The mystery of membrane organization: composition, regulation and roles of lipid rafts. Nat Rev Mol Cell Biol 18, 361– 374.

Sharonov, GV, Balatskaya, MN, and Tkachuk, VA (2016). Glycosylphosphatidylinositol-Anchored Proteins as Regulators of Cortical Cytoskeleton. Biochemistry (Mosc) 81, 636–650.

Simons, K, and Sampaio, JL (2011). Membrane organization and lipid rafts. Cold Spring Harb Perspect Biol 3, a004697.

Simons, K, and Toomre, D (2000). Lipid rafts and signal transduction. Nat Rev Mol Cell Biol 1, 31–39.

Slessareva, JE, Routt, SM, Temple, B, Bankaitis, VA, and Dohlman, HG (2006). Activation of the phosphatidylinositol 3-kinase Vps34 by a G protein alpha subunit at the endosome. Cell 126, 191–203.

Stringer, DK, and Piper, RC (2011). A single ubiquitin is sufficient for cargo protein entry into MVBs in the absence of ESCRT ubiquitination. Journal of Cell Biology 192, 229–242.

Suzuki, KGN, Kasai, RS, Hirosawa, KM, Nemoto, YL, Ishibashi, M, Miwa, Y, Fujiwara, TK, and Kusumi, A (2012). Transient GPI-anchored protein homodimers are units for raft organization and function. Nat Chem Biol 8, 774–783.

Szpurka, H, Schade, AE, Jankowska, AM, and Maciejewski, JP (2008). Altered lipid raft composition and defective cell death signal transduction in glycosylphosphatidylinositol anchor-deficient PIG-A mutant cells. Br J Haematol 142, 413–422.

Tatebayashi, K, Yamamoto, K, Nagoya, M, Takayama, T, Nishimura, A, Sakurai, M, Momma, T, and Saito, H (2015). Osmosensing and scaffolding functions of the oligomeric four-transmembrane domain osmosensor Sho1. Nat Commun 6, 6975.

Tognetti, S, Jiménez, J, Viganò, M, Duch, A, Queralt, E, de Nadal, E, and Posas, F (2020). Hog1 activation delays mitotic exit via phosphorylation of Net1. Proc Natl Acad Sci U S A 117, 8924– 8933.

Vazquez, HM, Vionnet, C, Roubaty, C, and Conzelmann, A (2014). Cdc1 removes the ethanolamine phosphate of the first mannose of GPI anchors and thereby facilitates the integration of GPI proteins into the yeast cell wall. Mol Biol Cell 25, 3375–3388.

Vogt, MS, Schmitz, GF, Varón Silva, D, Mösch, H-U, and Essen, L-O (2020). Structural base for the transfer of GPI-anchored glycoproteins into fungal cell walls. Proc Natl Acad Sci U S A 117, 22061–22067.

Yamamoto, A, DeWald, DB, Boronenkov, IV, Anderson, RA, Emr, SD, and Koshland, D (1995). Novel PI(4)P 5-kinase homologue, Fab1p, essential for normal vacuole function and morphology in yeast. Mol Biol Cell 6, 525–539.

Yang, G, and Banfield, DK (2020). Cdc1p is a Golgi-localized glycosylphosphatidylinositol-anchored protein remodelase. Mol Biol Cell 31, 2883–2891.

Yin, QY, de Groot, PWJ, Dekker, HL, de Jong, L, Klis, FM, and de Koster, CG (2005). Comprehensive proteomic analysis of Saccharomyces cerevisiae cell walls: identification of proteins covalently attached via glycosylphosphatidylinositol remnants or mild alkali-sensitive linkages. J Biol Chem 280, 20894–20901.

Yin, QY, de Groot, PWJ, de Jong, L, Klis, FM, and De Koster, CG (2007). Mass spectrometric quantitation of covalently bound cell wall proteins in Saccharomyces cerevisiae. FEMS Yeast Res 7, 887–896.

Zeng, L, Liu, H, Liu, Z, Li, L, Wang, H, Chen, Y, Wu, J, Wang, G, Li, L, and Fu, R (2023). Defected lipid rafts suppress cavin1-dependent IFN-α signaling endosome in paroxysmal nocturnal hemoglobinuria. Int Immunopharmacol 115, 109468.

Zhao, X, Li, R, Lu, C, Baluška, F, and Wan, Y (2015). Di-4-ANEPPDHQ, a fluorescent probe for the visualisation of membrane microdomains in living Arabidopsis thaliana cells. Plant Physiol Biochem 87, 53–60.

